# Comparative metabolomics identifies recurrent age-associated pathway remodeling across species

**DOI:** 10.64898/2026.06.23.733839

**Authors:** Genevieve A. Mortensen, Emily E. Montgomery, Innocent Ng’Ombwa, Shannon R. Smoot, Daniel Stephenson, Thomas C. Kaufman, Travis Nemkov, Angelo D’Alessandro, Laura M. Hurley, Jason M. Tennessen, Haixu Tang, Daniella E. Chusyd

**Affiliations:** Department of Computer Science, Indiana University, Bloomington, Indiana, USA; Department of Biology, Indiana University, Bloomington, Indiana, USA; Department of National Parks and Wildlife, Lusaka, Zambia; Department of Biochemistry and Molecular Genetics, University of Colorado Anschutz School of Medicine, Aurora, Colorado, USA; Department of Environmental and Occupational Health, Indiana University, Bloomington, Indiana USA

**Keywords:** Aging, Metabolomics, *Drosophila melanogaster*, Mice, Elephants, Fatty Acids, Carnitine

## Abstract

Aging is accompanied by widespread metabolic change, but it remains unclear which features are shared across species with different physiology, lifespan, and sampling contexts. To address this, we employed a pathway-centered comparative metabolomics framework to evaluate age-associated metabolic remodeling across wild African savanna elephant, mouse, and *Drosophila melanogaster. Drosophila* provided a controlled adult time course to map age-associated metabolite trajectories, while mouse and elephant plasma datasets allowed us to test whether these pathway signatures extended to mammalian aging. Adult *Drosophila* showed extensive metabolomic remodeling, with significant metabolites organizing into distinct temporal trajectory classes. Cross-species aging-associated metabolic correspondence was more apparent at the pathway level than at the level of individual metabolites. Pathway scores derived from *Drosophila* increased progressively with fly age, successfully distinguished young and old mice, and captured age-associated stratification across the elephant lifespan. Notably, lipid metabolism, particularly carnitine and fatty acid metabolism, together with nucleotide-related pathways, consistently emerged as the core features of aging across analyses. These findings suggest pathway-level metabolic remodeling is a recurrent feature of cross-species aging.

## INTRODUCTION

Aging is the major risk factor for many chronic diseases and is accompanied by progressive physiological decline across tissues and organ systems. At the molecular level, aging involves interconnected changes in nutrient sensing, mitochondrial function, proteostasis, inflammation, intercellular communication, and other processes that influence healthspan and disease vulnerability (López-Otín et al., 2023). Metabolism is central to these processes because it integrates gene regulation, nutrient availability, mitochondrial activity, environmental exposure, and physiological state. Age-associated metabolic remodeling may therefore provide a functional readout of biological aging that complements genomic, transcriptomic, epigenetic, and proteomic measures.

Given its high throughput and sensitivity, metabolomics has emerged as a powerful tool for mapping molecular signatures of aging across large cohorts. Human studies show that circulating metabolites are associated with chronological age, biological aging, mortality, longevity, frailty, chronic disease burden, and age-related disease risk (Huang et al., 2025; Ibáñez de Opakua et al., 2025; Kuiper et al., 2023; Menni et al., 2013; Mutz et al., 2024; Sebastiani et al., 2024, Johnson et al., 2019). Resources such as MetaboAge highlight the ubiquity of age-associated metabolic changes across diverse studies, although individual metabolite signatures vary by cohort, sample type, analytical platform, and study design (Bucaciuc Mracica et al., 2020). Recurring age-associated features include lipid metabolism, amino acid metabolism, nucleotide metabolism, mitochondrial and redox-related pathways, and central carbon metabolism (Huang et al., 2025, Reisz et al., 2025). While these findings support metabolomics as a strategy for studying aging, they also highlight a major challenge: individual metabolite signatures are often context-dependent and may not translate directly across species, tissues, or experimental designs.

This context dependence is especially important for cross-species aging studies. Blood-based proteomic and multiomic work has shown that molecular aging can vary by organ (Oh et al., 2023; Dzieciatkowska et al., 2026). Although focused on proteins rather than metabolites alone, they underscore that molecular aging signatures are unlikely to be uniform across biological matrices or organ systems. Pathway- and system-level analyses may therefore be more appropriate than individual molecular markers when translating age-associated signatures across organisms with different physiologies, sample types, and lifespans.

Model organisms have been essential for identifying conserved mechanisms of aging because they allow controlled manipulation of genetics, diet, environment, and adult age. *Drosophila melanogaster* is particularly useful because it is short-lived, genetically tractable, and amenable to dense adult time-course sampling (Piper and Partridge, 2018). It is also a well-established model for studying diet-sensitive aging mechanisms, nutrient-response pathways, and metabolic homeostasis (Evangelakou et al., 2019). Recent metabolomics studies in *Drosophila* show that adult aging is characterized by broad remodeling of metabolite abundance and metabolic activity, including changes in amino acid, lipid, nucleotide, and energy-associated pathways (Wang et al., 2022; Zhao et al., 2022, Vecchie et al., 2026). Stable-isotope tracing further indicates that aging disrupts coordinated metabolic flux across the fly metabolome, including glycolysis, serine metabolism, and purine metabolism (Wang et al., 2022). Adult fly studies have also linked high-sugar feeding to altered purine catabolism and shortened survival, connecting diet-sensitive metabolism, nucleotide turnover, and age-associated physiology (van Dam et al., 2020). Together, these findings support *Drosophila* as a time-resolved system for defining age-associated metabolic trajectories.

However, flies differ substantially from mammals in physiology, tissue organization, lifespan scale, and metabolite coverage. Thus, age-associated signatures discovered in fly studies are most useful for cross-species comparison when interpreted at the level of pathways or metabolic modules rather than as one-to-one corresponding metabolite markers. This approach is supported by systems-metabolism work in *Drosophila* showing that diet-induced metabolic dysregulation can be resolved at reaction and pathway levels using tissue-specific metabolic models and metabolomics-informed analyses (Moon et al., 2026). Such studies emphasize that while metabolite-level changes may vary across tissues or species, they often still converge on the same underlining biochemical processes.

Mammalian aging studies provide a complementary perspective. Mouse models allow controlled mammalian aging comparisons and have revealed widespread age-associated metabolic changes (Di Francesco et al., 2024, Jankowski et al., 2025). Long-lived mammals add another dimension because they may reveal metabolic features associated with aging across much longer lifespan scales. Comparative aging biology emphasizes that studies across diverse species can identify both shared and species-specific mechanisms of longevity, stress resistance, and disease susceptibility (Holtze et al., 2021; Ma & Gladyshev, 2017). Elephants are especially relevant because they are long-lived mammals with biological features that have motivated comparative studies of aging and disease resistance (Chusyd et al., 2021). However, metabolomic aging data from elephants and other long-lived non-traditional mammalian systems remain limited compared with data from humans, mice, and short-lived model organisms.

A central challenge in cross-species metabolomics is that direct metabolite-level comparisons are often constrained by differences in physiology, sample type, analytical coverage, annotation depth, and study design. As a result, the same biological process may be represented by different metabolites across species, even when similar age-associated metabolic remodeling is occurring. These differences can obscure shared biological patterns when analyses focus exclusively on individual metabolites. Pathway-centered approaches can address this limitation by evaluating whether age-associated metabolic changes converge on common biological processes, allowing common aspects of aging biology to be identified despite differences in the specific metabolites observed across species.

In this paper, we apply a *Drosophila*-anchored cross-species metabolomics framework to evaluate age-associated pathway remodeling across *Drosophila melanogaster*, mouse, and African savanna elephant *(Loxodonta africana)*. We first define age-associated metabolites and temporal trajectory classes in adult *Drosophila*, establishing the fly metabolome as a time-resolved reference. We then aggregate these signals to the pathway level and evaluate whether the same pathway categories show age-associated structure in mouse and African savanna elephant datasets. This design does not assume strict correspondence of individual metabolites. Instead, it tests whether a subset of *Drosophila* age-associated pathways exhibits broader correspondence across evolutionarily divergent species. By integrating a controlled invertebrate adult time course, a mammalian young-versus-old comparison, and a long-lived mammalian age gradient, this study identifies recurrent age-associated metabolic pathways and decouples shared pathway-level remodeling from species-specific metabolite profiles.

## METHODS

### Experimental design

This study compared age-associated metabolomic patterns across three species representing complementary lifespan scales and study designs: *Drosophila melanogaster* as a short-lived adult time-course model, *Mus musculus* as a laboratory mammalian young-versus-old comparison, and *Loxodonta africana* as a long-lived wild mammalian age-gradient dataset. The primary analysis was restricted to adult whole-fly samples for *Drosophila*, mouse plasma, and elephant plasma. Developmental stages, food controls, fecal samples, pooled samples, blanks, technical controls, serum samples, and mixed plasma/serum samples were excluded from the primary biological matrices; pooled samples, blanks, and technical controls were retained for quality-control assessment where available. Sample normalization, detection filtering, missing-value handling, and transformation were performed separately within each species as described below. The analytical framework was organized around *Drosophila* as the time-resolved anchor: age-associated metabolites and temporal trajectories were defined in flies, then evaluated at the pathway level in mouse and elephant.

### Drosophila husbandry

Oregon-R *Drosophila melanogaster* flies (RRID:BDSC_25211) were maintained on bottles of Semi-Defined food (https://bdsc.indiana.edu/information/recipes/germanfood.html) at 25°C with 50% relative humidity under a 12:12 h light:dark cycle. Following emergence of the first adult flies, ten independent stock bottles were cleared and flies allowed to eclose for five days, at which point 0–5-day old flies were separated by sex. At this point, a 0-5 day sample of 20 males and 20 females were collected from each bottle and the remaining flies were transferred to vials containing freshly prepared semi-defined food for subsequent collections at 12-18 days and 25-30 days. We encode each timepoint in our analyses using the oldest day in each range.

Vials were kept at 25°C with 50% relative humidity under a 12 h light/12 h dark cycle and transferred to new food every 2-3 days without CO2 anesthesia. Flies were transferred to empty vials, immobilized on dry ice, counted, transferred to tared bead-containing tubes, weighed, snap-frozen in liquid nitrogen, and stored at −80°C as previously described (Holsopple et al., 2023). Frozen samples were subsequently prepared for liquid chromatography–mass spectrometry using a published protocol (Fleck et al., 2024). FlyBase was used as a genetic resource (Öztürk-çolak et al., 2024).

### Mouse housing

Mouse plasma samples were obtained from a previously conducted socialization study in CBA/J mice. The analyzed subset comprised 20 mice, including four young males (4–5 months of age) housed under social conditions, four old males and four old females housed under social conditions, and four old males and four old females housed under isolation conditions (12-18 months of age). Mice were maintained under standard laboratory conditions with ad libitum access to food and water under a 14:10 h light:dark cycle. Plasma was collected into 0.2 mL microcentrifuge tubes following centrifugation and stored at −80°C. For the present comparative aging analysis, age was the primary variable of interest, while sex and housing condition were included as covariates in the metabolite-level models. Because the experimental design did not include young females or young isolated mice, age effects were interpreted within the structure of the available groups rather than as fully factorial age-by-sex or age-by-housing effects.

### Wild elephant sampling

African savanna elephant plasma samples were collected in collaboration with Game Rangers International, Zambia’s Department of National Parks and Wildlife, and African Parks as part of a broader study of orphaned and non-orphaned elephants in Kafue National Park, Zambia. Between May 2022 and June 2024, whole blood was collected from anesthetized elephants during translocation or GPS collaring procedures. The dataset included 23 individuals, 12 females and 11 males, aged 2–23 years, with two males sampled twice one year apart. Two replicate plasma samples were collected from each elephant at each of 25 sampling events, yielding 50 total plasma samples. Blood was collected into EDTA tubes, centrifuged within 60 min, aliquoted, stored at −20°C in Zambia, shipped on ice to Indiana University, and stored at −80°C. Age was known within 3 months for most individuals or estimated from shoulder height; sex was recorded from external genitalia (Hanks, 1972).

### Sample preparation and metabolomics

Frozen samples were extracted with prechilled 90% methanol containing 2 μg/mL succinic-d4 acid. Samples were homogenized, incubated at −20°C for 2 h, centrifuged, and supernatants were transferred, re-centrifuged, aliquoted into 96-well plates, dried under nitrogen, and stored at −80°C. Ultra-high-pressure liquid chromatography–mass spectrometry metabolomics was performed at the University of Colorado Anschutz Medical Campus as previously described (Nemkov et al., 2019). Extracts were analyzed using a Vanquish ultra-high-pressure liquid chromatography system coupled to a Q Exactive mass spectrometer operated in positive and negative ion modes. Metabolites were resolved on a Kinetex C18 column using polarity-specific mobile phases, and full-scan data were acquired from 60 to 900 m/z at 70,000 resolution.

### Data preprocessing

Preprocessing and statistical analyses were performed in Python, with analysis code available at https://github.com/ginnymortensen/Aging_Metabolome.git. ChatGPT Codex was used to assist with script troubleshooting and code debugging. The metabolomics workbooks contained quantitative LC–MS feature-abundance measurements for individual annotated metabolites. These values represent the analytical signal reported for each metabolite feature rather than transformed or standardized statistical values. The raw matrices contained 290 metabolite features for *Drosophila* and 343 metabolite features for each mammalian dataset.

Samples were first classified according to biological sample type and study design. Adult whole-fly samples, mouse plasma samples, and elephant plasma samples were retained for the primary comparative analyses. Developmental-stage fly samples, mouse and elephant fecal samples, pooled samples, blanks, other technical controls, elephant serum samples, and mixed plasma/serum samples were excluded from the primary biological matrices. Pooled samples, blanks, and other controls were retained during preprocessing for quality-control assessment where available but were not included as biological observations in downstream age-associated analyses. Because the mammalian datasets were acquired on separate experimental plates, analyses were performed within species rather than by combining raw intensities across species.

Zero values were interpreted as undetected or non-quantifiable measurements and were treated as missing values before logarithmic transformation. No negative metabolite-abundance values were present in the analyzed raw matrices. Consequently, no negative values were generated or subsequently required to be removed because of the normalization procedure described below.

For *Drosophila*, metabolite abundances were normalized to sample mass prior to sample-level scaling. For mouse and elephant plasma, abundances were normalized to the recorded amount of material used for extraction. Following material normalization, a sample-median normalization procedure was applied. Specifically, for each biological sample, the median abundance across all measured metabolite features was calculated. The reference value for normalization was then defined as the median of these sample-specific medians across all biological samples in the corresponding species dataset. Each metabolite abundance within a sample was multiplied by the ratio of the reference value to that sample’s median abundance. Thus, “global median” refers to the median across samples of the sample-specific metabolite medians; it does not refer to the median abundance of metabolites within an individual sample. Because this is a multiplicative scaling procedure, it does not generate negative abundance values.

Metabolites were retained when detectable in at least 70% of biological samples within the corresponding species dataset. After filtering, remaining missing values were imputed separately for each metabolite using one-half of the minimum positive observed abundance for that metabolite. The resulting complete abundance matrices were log2 transformed for statistical modeling. Feature-wise z-score transformation of the log2-transformed data was used only for visualization and cross-metabolite comparisons and was not used to generate metabolite-level statistical tests. Near-zero-variance features were removed after log2 transformation. The final matrices contained 250 adult *Drosophila* metabolite features, 254 mouse plasma metabolite features, and 225 elephant plasma metabolite features.

Technical variation was considered separately from biological variation. Sample mass or material amount was incorporated into the normalization procedure, while pooled samples, blanks, and technical controls were used for quality-control assessment rather than treated as biological replicates. Because the available pooled-control structure did not provide repeated matrix-matched quality-control injections distributed throughout each analytical run, formal signal-drift correction was not applied. Mouse and elephant datasets were analyzed within their respective analytical runs, and differences in sample type, experimental plate, and study design were not interpreted as biological effects. Internal-standard measurements were monitored as indicators of analytical performance but were not used as a universal normalization factor because the internal standard was not established as an appropriate proportional correction for all measured metabolites.

### Age-effect modeling

Age-associated metabolites were identified separately within each species using metabolite-wise regression models fit to log2-transformed metabolite abundances. For each metabolite, the age effect was estimated as the coefficient of the primary age term while adjusting for available covariates.

In *Drosophila*, age was modeled as a standardized continuous adult-age variable, with sex included as a covariate:

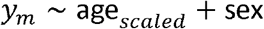

In mouse, age was modeled as a binary old-versus-young variable, with sex and social condition (isolated or social) included as covariates:

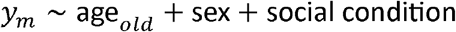

In elephant, age was modeled as a standardized continuous age variable, with sex included as a covariate and individual identity included as a random intercept to account for repeated sampling:

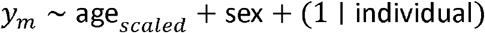

Ordinary least-squares models were used for *Drosophila* and mouse. Elephant models were fit using linear mixed-effects models with a random intercept for individual; if mixed-model fitting failed, an ordinary least-squares model with cluster-robust standard errors by individual was used as a fallback.

For each metabolite, the age-effect coefficient, standard error, test statistic, and t-test computed p-value were extracted from the primary age term. P-values were adjusted within each species using the Benjamini–Hochberg false discovery rate (FDR) procedure. Metabolites with positive age coefficients were classified as age-increasing, and metabolites with negative coefficients were classified as age-decreasing.

### Pathway mapping and overlap analyses

Metabolites were assigned identifiers from the Human Metabolome Database (HMDB) or Kyoto Encyclopedia of Genes and Genomes (KEGG) and mapped to curated metabolic pathways. The targeted mammalian panel contained 343 analytical metabolite features, including multiple analytical ions corresponding to the same compound. Across the panel, these features were assigned to 41 curated pathway categories, representing 324 unique metabolite– pathway relationships.

Pathway overlap was evaluated using complementary descriptive approaches designed to account for differences in statistical power and metabolite coverage among species. First, age-associated metabolites were ranked within each species according to the (p)-value from the species-specific age-effect model, with smaller (p)-values indicating stronger evidence for an association with age. Because the mouse and elephant datasets did not yield FDR-significant individual metabolites at (q < 0.05), ranked analyses were used to characterize patterns among the strongest age-associated features without treating a nominal threshold as evidence of statistical significance. For the matched-size comparison, the number of mouse metabolites meeting a nominal (p < 0.10) threshold (35 metabolites) was used solely to define a common feature count across species. The 35 highest-ranked age-associated metabolites from each species were therefore compared for pathway representation. The nominal threshold criterion was not used to define FDR-significant metabolites, and results from this analysis were interpreted as exploratory pathway-overlap patterns rather than formal evidence of age-associated pathway enrichment.

To assess the sensitivity of this ranked comparison to the choice of feature count, pathway representation was additionally examined across alternative top-(N) ranked metabolite sets. These analyses were used to determine whether recurrent pathway categories were apparent across a range of ranked feature counts rather than being dependent on a single threshold or feature number.

Second, a *Drosophila*-anchored overlap analysis was performed. FDR-significant *Drosophila* age-associated metabolites (q < 0.05) were mapped to pathways and compared with metabolites that were detected and eligible for statistical modeling in mouse and elephant. This analysis asked whether the biological pathways represented among robust *Drosophila* age-associated metabolites were also represented in the mammalian datasets, even when the exact metabolite was not measured or did not show a statistically significant age association in the mammalian data.

These overlap analyses were intended to describe the extent and composition of pathway correspondence across species rather than to provide an independent statistical test of pathway conservation. Because pathway size, metabolite coverage, metabolite annotation, and the correlation structure of metabolites differ among pathways and species, simple pathway overlap was not interpreted as evidence that the observed correspondence exceeded a defined random expectation. Formal age-associated effects were evaluated separately at the individual-metabolite level using species-specific statistical models and FDR correction.

### Global aging scoring

A *Drosophila*-anchored pathway metabolic aging score was calculated to summarize whether pathways associated with aging in *Drosophila* showed corresponding age-related metabolic structure in mouse and elephant. First, sex-adjusted metabolite-level models were used to identify FDR-significant age-associated metabolites in *Drosophila* using an FDR threshold of 0.05. These metabolites were mapped to annotated pathways, and pathways containing at least one FDR-significant *Drosophila* metabolite were retained as *Drosophila*-associated aging pathways. This criterion was used to define the pathway set represented among the strongest *Drosophila* metabolite-level age associations and was not intended as an independent pathway-level significance test.

Each retained pathway was then scored separately within each species using that species’ own detected and modelable metabolites assigned to the same pathway. Here, “available” refers to metabolites that were detected in the corresponding dataset, passed the metabolite-level filtering criteria, and could be included in the species-specific age-effect model. Metabolites were classified as age-increasing or age-decreasing according to the sign of the species-specific age-effect coefficient. Specifically, for sample *i* and pathway *p*, the pathway score was calculated as:

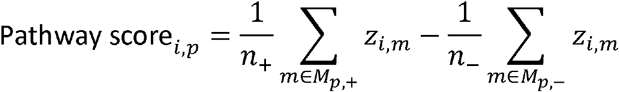

where *Z*_*i*,*m*_is the feature-wise z-scored abundance of metabolite *m* in sample *i, M*_*p*,+_and *M*_*p*,-_ denote the age-increasing and age-decreasing metabolites assigned to pathway *p*, and *n*_+_and *n*_-_are the corresponding metabolite counts. Thus, metabolites with positive age effects contribute positively to the pathway score, whereas metabolites with negative age effects are directionally reversed. Pathways were scored only when at least two species-specific metabolites were available.

The global pathway aging score for each sample was then defined as the mean of the available pathway scores:

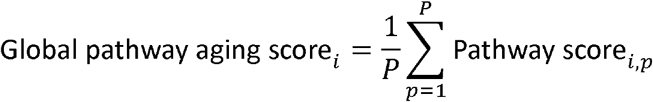

where *P* is the number of retained *Drosophila*-anchored pathways included in the cross-species analysis. For cross-species comparisons, only pathways that could be scored in all three species were retained so that the same predefined pathway set contributed to each species-level summary; this was an availability criterion rather than a statistical significance criterion.

The pathway aging score was used as a composite summary of age-associated metabolic structure. Because the pathway set and metabolite directions were derived from the species-specific age-effect analyses, the score was not treated as an independent test of pathway conservation. Statistical evidence for individual metabolite age associations was established separately using the species-specific regression models and FDR correction described above.

### Trajectory clustering and concordance scoring

*Drosophila* age-associated metabolites were defined from sex-adjusted age-effect models using q < 0.05. For each significant metabolite, mean z-scored abundance was calculated across the three adult age groups. The resulting metabolite-by-age matrix was row-scaled so that clustering reflected relative temporal pattern rather than absolute abundance. Metabolites were grouped into five trajectory classes using k-means clustering with 50 random initializations and a fixed random seed. Clusters were ordered by net change between 5 and 30 days, and centroids were used to summarize dominant temporal patterns.

To evaluate whether *Drosophila* trajectory-associated pathways showed concordant age effects in mammals, metabolites within each fly trajectory cluster were mapped to pathways. For each trajectory cluster–pathway pair, mouse and elephant age-effect coefficients were extracted for all tested metabolites assigned to the same pathway. Concordance was calculated as the product of the signed *Drosophila* trajectory direction and the mean species-specific age-effect coefficient for all tested metabolites in that pathway. Positive values indicate mammalian age effects concordant with the *Drosophila* trajectory direction, whereas negative values indicate opposing directionality.

## RESULTS

### Cross-species metabolomics framework for evaluating shared age-associated pathway remodeling

To evaluate whether aging is associated with shared metabolic remodeling across evolutionarily divergent species, we integrated metabolomics datasets from *Drosophila melanogaster*, mouse, and African savanna elephant (Fig. 1). These species span markedly different lifespan scales and experimental contexts, with *Drosophila* providing a controlled adult time course, mouse providing a mammalian young-versus-old contrast, and elephant providing a cross-sectional age gradient in a long-lived mammal. Following targeted LC/MS metabolomics, all datasets were processed through a standardized workflow including normalization, metabolite detection filtering, missing-value imputation, log2 transformation, and feature-wise scaling. The analytical strategy was structured around a *Drosophila*-anchored framework: age-associated metabolites and temporal trajectory classes were first defined in flies, then aggregated to the pathway level, and finally evaluated in mouse and elephant to test whether fly-derived age-associated pathways showed evidence of age-related remodeling across mammals. Additional supporting analyses were used to evaluate sex-specific structure, pathway-overlap sensitivity, score robustness, metabolite directionality, and trajectory-concordance coverage across species (Supplementary Figs. S1–S13).

**Figure 1.**
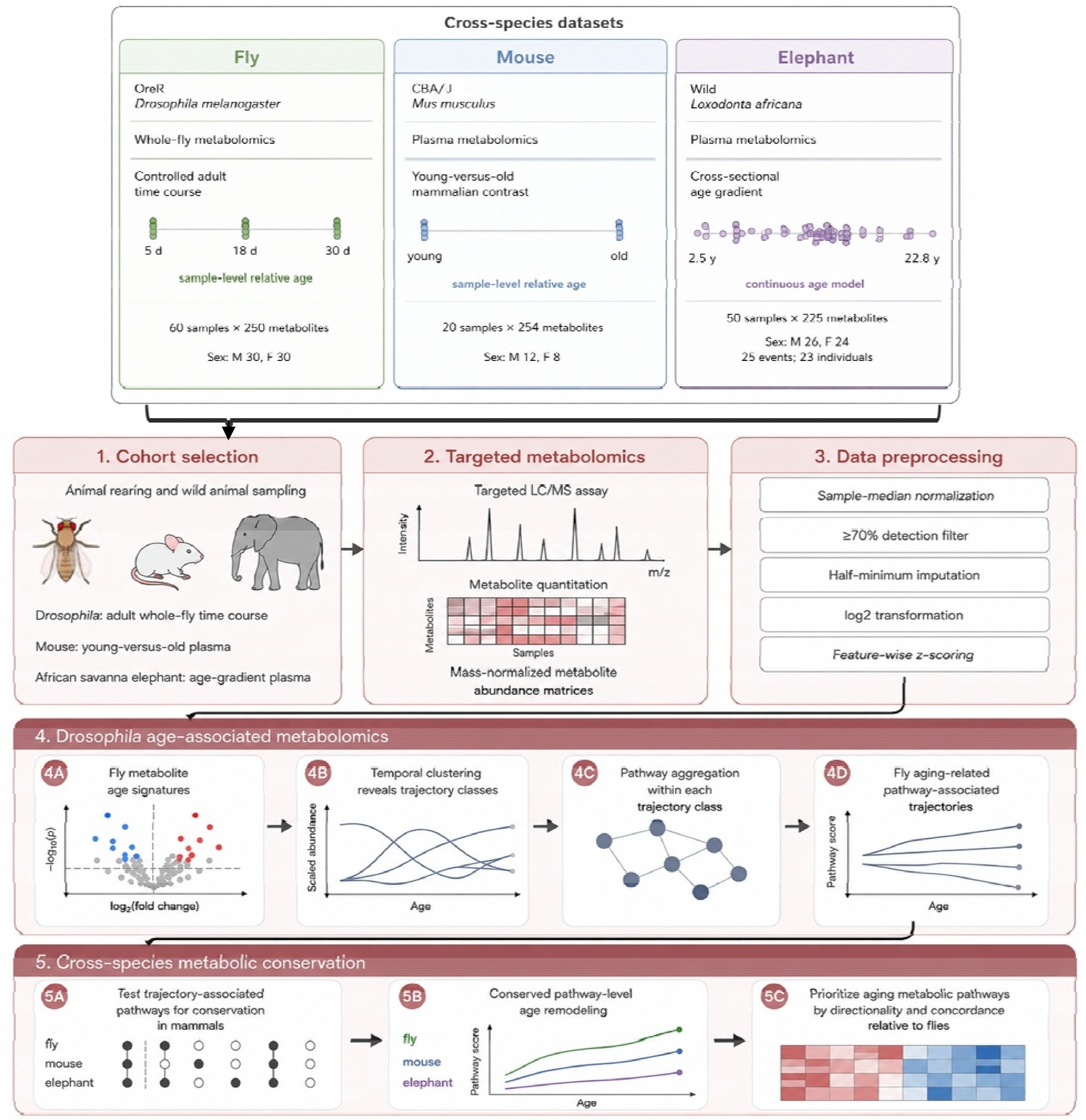
Cross-species metabolomics study design and computational workflow. The study compares age-associated metabolomic patterns across Drosophila melanogaster, mouse, and African savanna elephant. Drosophila whole-fly samples provide a controlled adult time course, mouse plasma samples provide a young-versus-old mammalian comparison, and elephant plasma samples provide a cross-sectional age gradient. Samples were analyzed using targeted LC/MS metabolomics, followed by sample-median normalization, metabolite detection filtering, half-minimum imputation, log2 transformation, and feature-wise z-scoring. Computational analyses were performed in a Drosophila-anchored framework: age-associated metabolites were identified in flies, grouped into temporal trajectory classes, aggregated to pathways, and then tested for pathway-level age remodeling in mouse and elephant. Metabolite-level age associations were evaluated using species-specific statistical models with FDR correction; pathway-overlap, score-based, and trajectory analyses were used primarily to summarize biological structure and were interpreted as descriptive or exploratory unless an independent statistical test is specified.

### Adult aging is associated with broad metabolomic remodeling in *Drosophila*

Principal component analysis of the *Drosophila* metabolomics dataset showed clear separation of samples by adult age, indicating that chronological age was a dominant source of metabolomic variation (PC1 = 53.9%) while additional structure associated with sex (PC2 = 11.7%) (Fig. 2A). To identify individual metabolites associated with adult age, sex-adjusted metabolite-level models were fit for each measured metabolite, testing the null hypothesis that metabolite abundance was not associated with age after adjustment for sex. Of the 223 metabolites showing nominal evidence of association (p < 0.05), 220 remained significant after Benjamini–Hochberg FDR correction (q < 0.05). Among these, asymmetric dimethyl arginine, acylcarnitine (6:0), hypoxanthine, N-acetyl-L-histidine, L-carnitine, and 5-hydroxyisourate decreased most significantly with age. Cyclic CMP, 10-formyltetrahydrofolate, 2-methyleneglutarate, and indole-3-acetate most significantly increased with age (Fig. 2B). Because the *Drosophila* dataset included both males and females across all age groups, supporting analyses further evaluated whether age-associated trajectory patterns contained sex-specific structure (Supplementary Figs. S1–S3).

**Figure 2.**
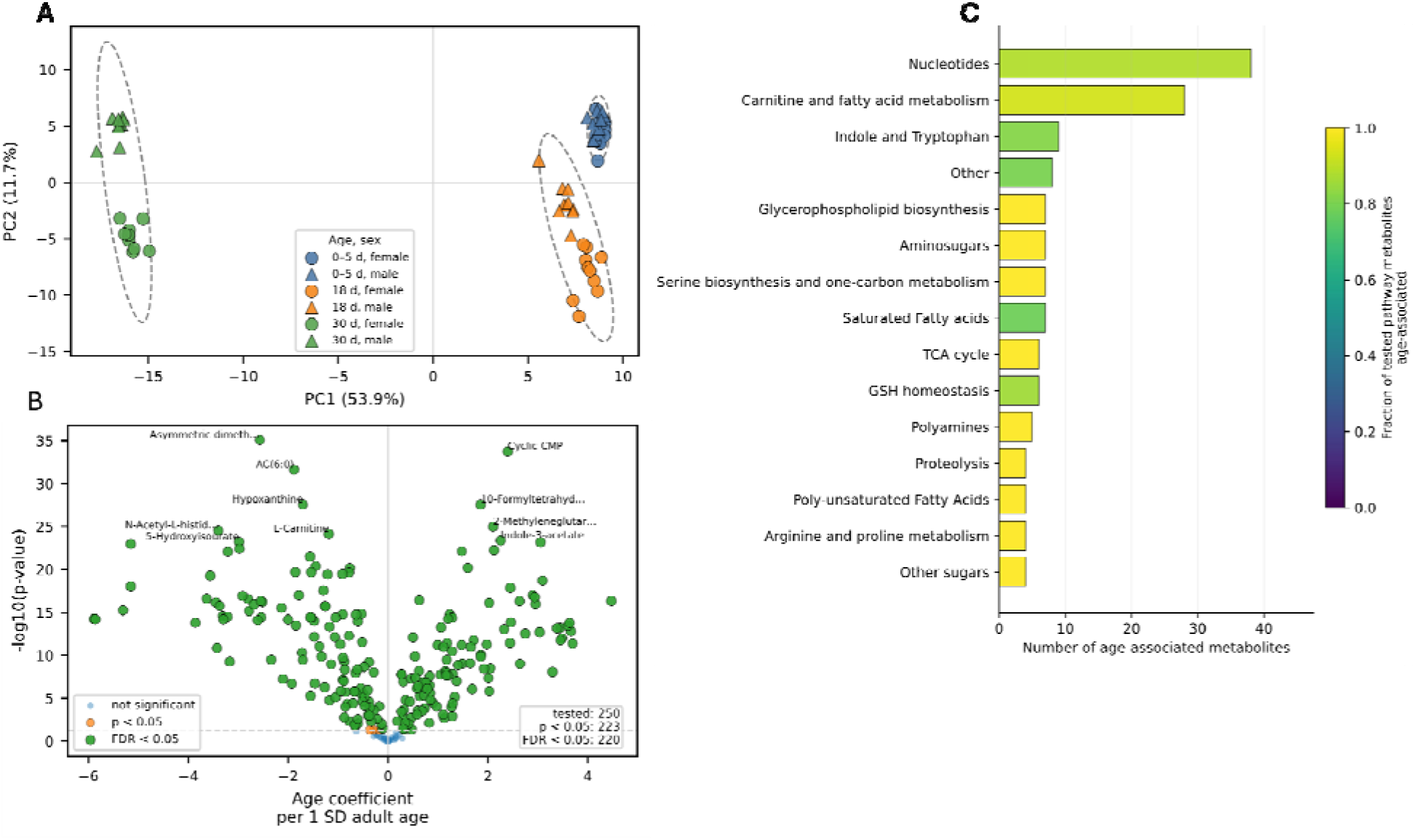
Adult aging is associated with extensive metabolomic remodeling in Drosophila. A: PCA of normalized Drosophila metabolite abundances, colored by age and shaped by sex. Percent variance contributed by each principal component is indicated in respective axes. B: Volcano plot of sex-adjusted metabolite-level age effects. For each metabolite, the null hypothesis was that abundance was not associated with age after adjustment for sex. The x-axis shows the estimated age coefficient, and the y-axis shows the nominal -log_10_(p). Metabolites meeting the FDR threshold (q < 0.05) are highlighted. C: Pathway representation of FDR-significant Drosophila age-associated metabolites. Bars show the number of FDR-significant metabolites assigned to each annotated pathway, and the color indicates the fraction of eligible metabolites within that pathway that were age-associated. This panel summarizes pathway representation among significant metabolites and is descriptive.

Pathway-level annotations of age-associated metabolites showed that several metabolic categories were prominently represented among the fly aging signatures. Age-associated metabolites were highly represented in nucleotides, carnitine/fatty acid metabolism, and indole/tryptophan metabolism (Fig. 2C). However, glycerophospholipid biosynthesis, amino sugar metabolism, serine biosynthesis and one-carbon-related metabolism, TCA cycle, and other amino acid related metabolism classes were exclusively represented by age-associated metabolites. Interestingly, many fatty acid classes were consistently highly represented as well. This indicates that aging in *Drosophila* is not characterized by a uniform global increase or decrease in metabolite abundance, but rather by coordinated remodeling across multiple diverse metabolic classes. These results echo prior studies using *Drosophila* as a time-resolved reference for defining age-associated metabolite signatures and pathway-level remodeling (Avanesov et al., 2014; Hoffman et al., 2014).

### Mouse and elephant datasets show age-associated metabolite signals with broader pathway-level correspondence

Age-associated metabolite modeling was performed next in the mammalian datasets. In mouse, sex and socialization-adjusted metabolite-level models were fit for each measured metabolite, testing the null hypothesis that metabolite abundance was not associated with age after adjustment for sex or socialization condition. In mouse, comparison of old versus young samples identified 21 metabolites with nominal evidence of age-association (p < 0.05); none remained significant after FDR correction (Fig. 3A). Because the mammalian datasets had substantially smaller sample sizes and less dense age sampling than the *Drosophila* time course, we also examined the strongest age-ranked features in an exploratory analysis. In mouse, 35 metabolites showed nominal age-association at p < 0.10, with most decreasing with age, including purine-related metabolites such as xanthine, hypoxanthine, guanosine, and inosine. FA(18:1) was among the metabolites that increased with age, and this set was used in corresponding overlap analysis (Fig. 3C & D).

**Figure 3.**
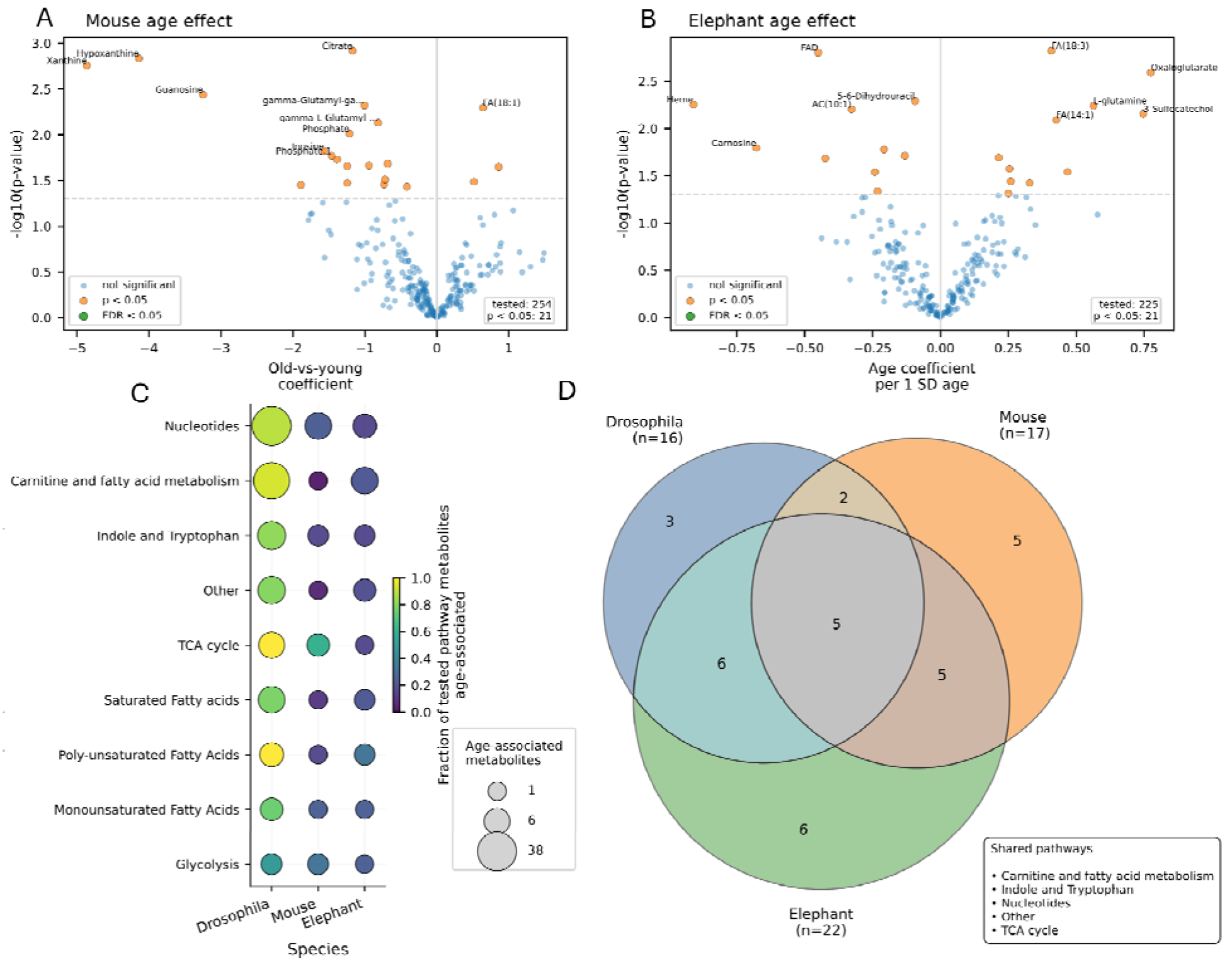
Age-associated metabolite and pathway signals in mouse and elephant. A: Volcano plot of the mouse old-versus-young metabolite comparison. Points indicate individual metabolites, with nominally significant age-associated metabolites highlighted. B: Volcano plot of continuous age effects in elephant plasma metabolites. Points indicate individual metabolites, with nominally significant age-associated metabolites highlighted. C: Descriptive comparison of pathway representation among age-associated metabolites across Drosophila, mouse, and African savanna elephant. Bubble size indicates the number of age-associated metabolites assigned to each pathway, and color indicates the fraction of eligible metabolites within that pathway that met the corresponding nominal age-association criterion. D: Venn diagram showing the observed overlap in annotated pathways represented among age-associated metabolites across species. The diagram summarizes pathway annotation overlap rather than a statistical test of whether the observed overlap exceeds a random expectation.

In African savanna elephants, sex-adjusted continuous age modeling of each measured metabolite tested the null hypothesis that metabolite abundance was not associated with age after adjustment for sex and accounting for repeated sampling. This modeling identified 21 metabolites with nominal evidence of age association (p < 0.05); none remained significant after FDR correction (Fig. 3B). To descriptively examine overlap correspondence across species, at (p < 0.10), 38 metabolites showed nominal age association and were used to characterize the strongest age-related metabolite signals across the sampled age gradient (Fig. 3C & D). Elephant age-associated metabolites included lipid, amino acid, redox, and nucleotide-related features, with several fatty acid metabolites increasing with age. Additional elephant analyses stratified age-associated metabolite quantity and effect by sex to support interpretation of age-related patterns in this cross-sectional dataset (Supplementary Fig. S4). Species-specific summaries of significant age-effect metabolites are provided in Supplementary Fig. S5.

Although the number and strength of individual metabolite-level associations were more limited in the mammalian datasets than in the *Drosophila* time course, both datasets showed evidence of age-associated metabolic variation. Because exact metabolite-level overlap across species may be constrained by species-specific physiology, sample type, and metabolite coverage, we next evaluated age-associated overlap at the pathway level (Fig. 3C). This analysis revealed that several pathways represented among *Drosophila* age-associated metabolites were also represented among nominally age-associated metabolites in mouse and/or elephant. These included lipid, nucleotide, amino acid, and central metabolism-related categories. Metabolite-to-pathway mapping of significant age-associated features is shown in Supplementary Fig. S6.

Because the mammalian datasets did not yield FDR-significant individual metabolites, we next examined the strongest age-associated features using a ranked analysis with a nominal threshold (p < 0.10) from age-effect modeling. 35 mouse metabolites met the nominal criterion, and this number was used solely to establish a matched feature count for the exploratory cross-species ranked comparison. The 35 highest-ranked age-associated metabolites from each species were therefore evaluated for pathway representation. Several pathway categories recurred across species, including carnitine and fatty acid metabolism, fatty acid subclasses, nucleotide metabolism, and central metabolic pathways (Fig. 3D). Examination across alternative ranked feature counts showed that several of these pathway categories remained represented across a range of top-(N) selections (Supplementary Fig. S7). These supplemental analyses were descriptive and were used to characterize pathway-level correspondence among the species rather than to establish statistically significant pathway conservation. Jaccard similarity analysis of whole age-associated pathway profiles provided an additional quantitative summary of pathway-level overlap across species (Supplementary Fig. S8). These findings support pathway-level comparison to evaluate cross-species age-associated structure while avoiding overinterpretation of one-to-one metabolite correspondence.

### *Drosophila*-anchored pathway aging scores reveal cross-species age-associated structure

To evaluate whether pathways associated with aging in *Drosophila* captured corresponding age-related metabolic structure in the mammalian datasets, we calculated pathway aging scores using 19 *Drosophila* age-associated pathways with sufficient metabolite coverage for scoring across all three species (Fig. 4A, Supplementary Table S1). In *Drosophila*, these scores increased across adult age, consistent with the age-associated metabolite patterns used to define the pathway set. In mouse, the same pathway-based scoring framework showed lower scores for young samples and higher scores for old samples, supporting an increasing trend as well. Elephant pathway scores also showed an increasing age-associated trend across the sampled chronological age gradient. Because the pathway set and metabolite directions contributing to the scores were derived from the species-specific age-effect analyses, these scores were interpreted as summaries of age-associated metabolic structure rather than independent statistical tests of pathway conservation. However, pathway-stratified scores are shown in Supplementary Fig. S10, allowing assessment of which individual pathways contributed to the global score structure in each species.

**Figure 4.**
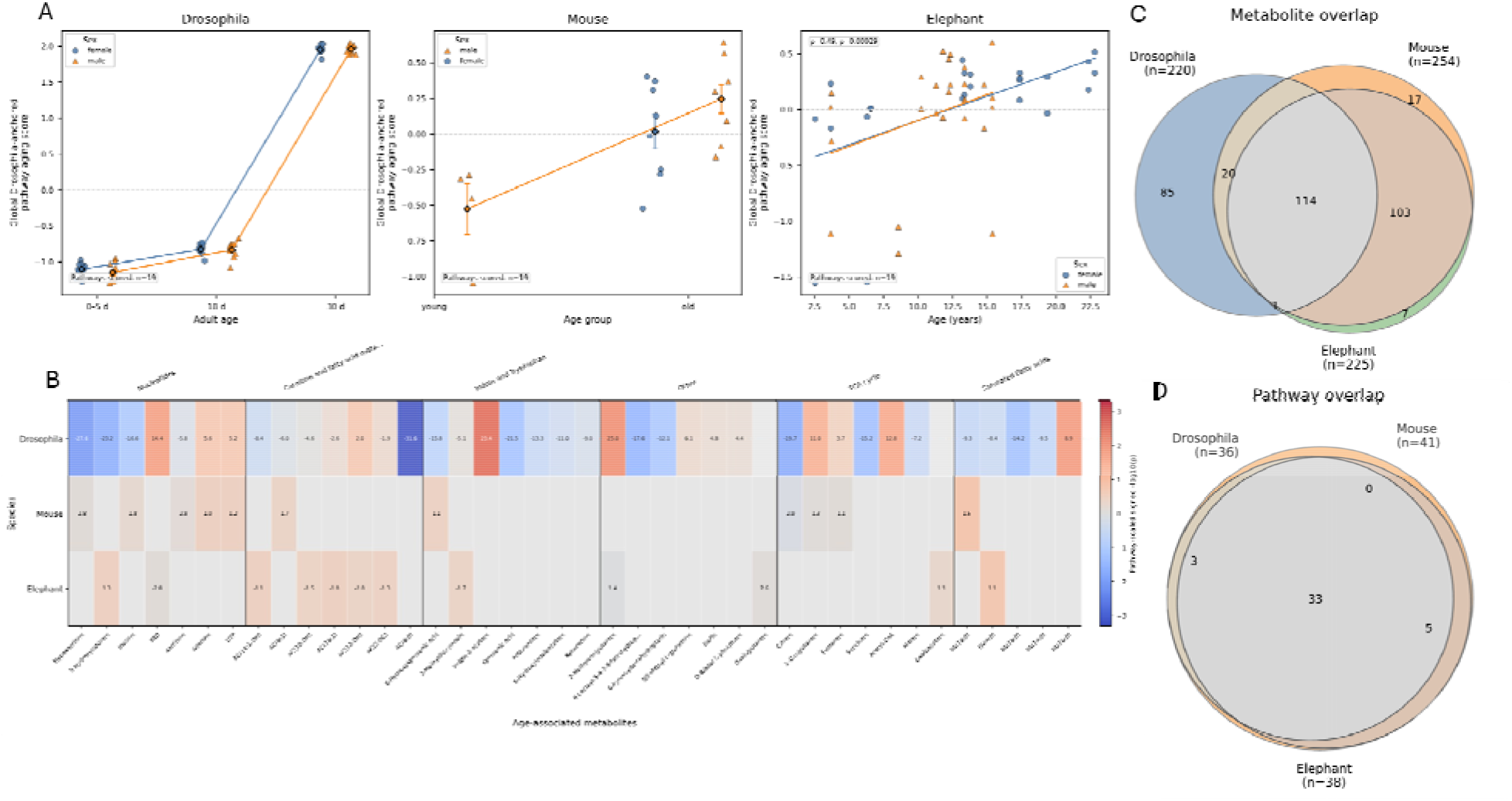
Drosophila-anchored pathway metabolic aging scores reveal cross-species age-associated remodeling. A: Global pathway-anchored metabolic aging scores for each species calculated from 19 Drosophila age-associated pathways. B: Heatmap of age-associated metabolites within shared pathways across species. Values represent pathway-scaled signed age effects. Gray cells indicate no metabolite-specific cross-species signal. C: Venn diagram showing the observed overlap in measured metabolite identities among Drosophila, mouse, and African savanna elephant. This panel describes analytical coverage and shared metabolite identity. D: Observed overlap in annotated metabolic pathways among Drosophila, mouse, and African savanna elephant. This panel describes annotation overlap.

Metabolite-level heatmap analysis further showed that shared pathways were composed of partially overlapping metabolite sets across species (Fig. 4B). Notably, many acylcarnitine and fatty acid variants are seen to accumulate with age in mammalian models. Direct overlap and UpSet analyses confirmed that cross-species correspondence was broader at the pathway level than at the individual-metabolite level (Fig. 4C and 4D, Supplementary Fig. S12). Directionality analyses also showed that pathways differed in the balance of age-increasing and age-decreasing metabolites across species (Supplementary Figs. S9 and S11). These results support a pathway-centered interpretation in which a subset of *Drosophila* age-associated pathways shows corresponding age-related remodeling in mouse and elephant.

### Drosophila age-associated metabolites resolve into distinct temporal trajectory classes

To determine whether fly age-associated metabolites followed common temporal patterns, FDR-significant Drosophila age-associated metabolites were clustered according to their abundance trajectories across adult age (Fig. 5A). The resulting clusters included metabolites showing progressive age-associated increases, progressive decreases, and non-monotonic or transient midlife-associated patterns. Cluster centroid trajectories confirmed that metabolites within each class shared similar temporal profiles, while representative metabolites demonstrated that each cluster contained biologically diverse features (Fig. 5B). Representative metabolites were selected as the lowest FDR with closest proximity to the respective cluster’s centroid. Because sex was an additional source of structure in the *Drosophila* dataset, supplemental analyses evaluated sex-specific metabolite counts, median female-minus-male differences, and root mean-squared sex-divergence scores within trajectory clusters (Supplementary Figs. S1–S3).

**Figure 5.**
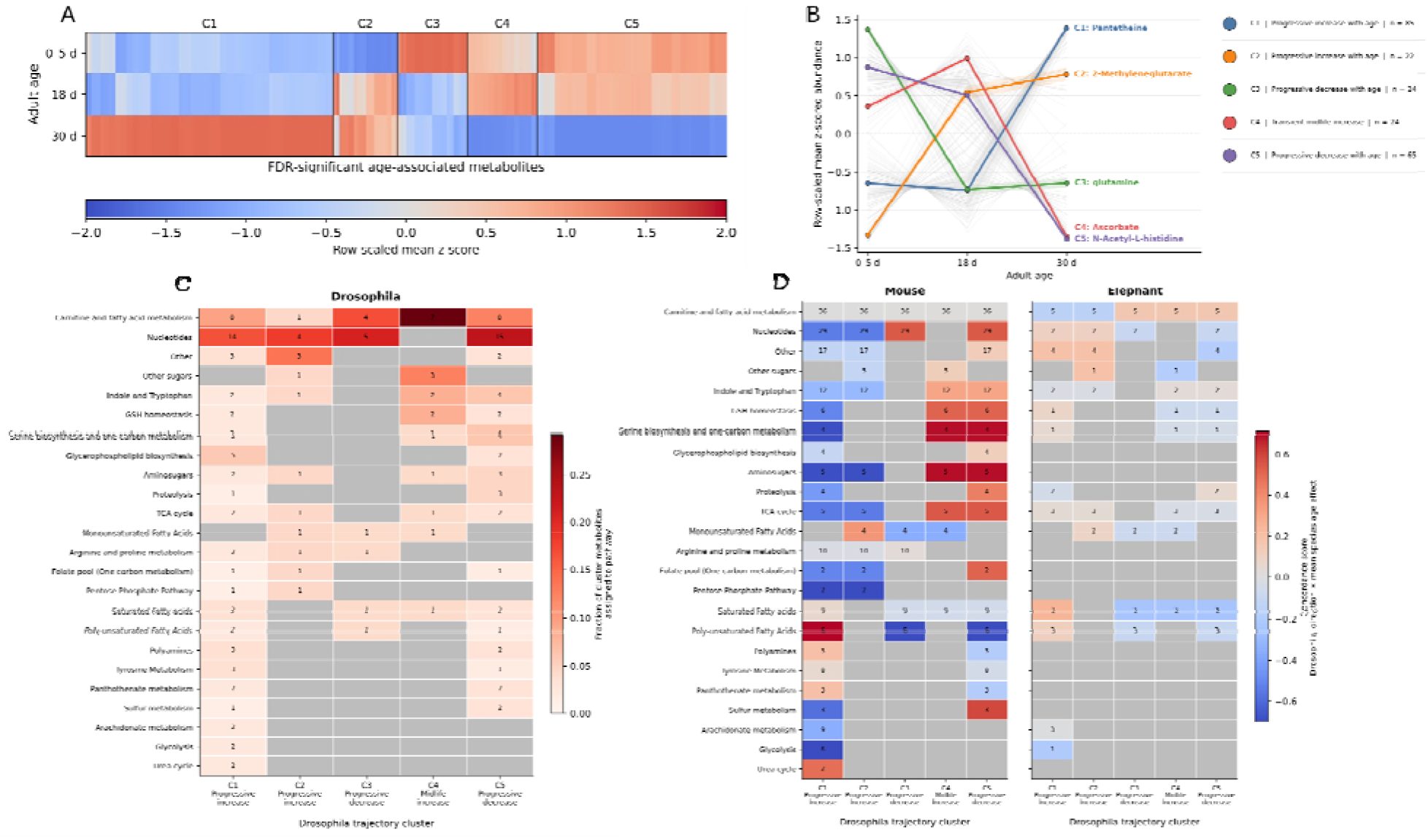
Temporal clustering of Drosophila age-associated metabolites identifies trajectory classes with pathway-level cross-species correspondence. A: Heatmap of FDR-significant Drosophila age-associated metabolites grouped by temporal trajectory cluster. Values represent row-scaled mean z-scored abundance across adult age groups. B: Cluster centroid trajectories showing representative temporal patterns for each metabolite cluster. Thin gray lines represent individual metabolite trajectories, while colored lines indicate cluster-level mean trajectories and labeled representative metabolites. C: Distribution of annotated metabolic pathways across Drosophila temporal trajectory clusters. Heatmap values indicate the fraction of metabolites within each cluster assigned to each pathway, with cell labels indicating metabolite counts. This panel describes pathway representation within the trajectory clusters. D: Cross-species evaluation of Drosophila trajectory-associated pathways in mouse and African savanna elephant. Heatmap values summarize the direction and magnitude of mammalian age effects for metabolites assigned to pathways represented within each Drosophila trajectory class. Positive values indicate concordance with the overall Drosophila trajectory direction, whereas negative values indicate opposing directionality. Gray cells indicate insufficient metabolite representation.

Pathway annotations were distributed differently across the Drosophila trajectory clusters, with nucleotide metabolism represented across multiple temporal patterns and carnitine/fatty acid metabolism particularly represented among metabolites showing transient midlife increases (Fig. 5C). Nucleotide metabolism was prominent across multiple trajectory classes, suggesting that this pathway undergoes complex age-associated remodeling rather than a single directional shift. However, carnitine and fatty acid metabolism showed especially strong influence in the transient midlife increase cluster trajectory. All fatty acid classes showed signal as progressively decreasing with age. Other pathways showed more cluster-specific representation, such as glycolysis and urea cycle in progressively increasing clusters, indicating that distinct metabolic processes may be remodeled at different phases of adult aging. These analyses identified temporal patterns among *Drosophila* age-associated metabolites which can be used to define trajectory-associated pathway signatures. Trajectory clustering was used to characterize similarity in temporal abundance profiles among metabolites already identified as age-associated and was therefore not an independent test of the null hypothesis of no age effect. Evidence for age association was established by the metabolite-level models described above, while clustering was used to summarize heterogeneity in the temporal patterns of those metabolites.

### Trajectory-associated *Drosophila* pathways show partial mammalian correspondence

Finally, *Drosophila* trajectory-associated pathways were evaluated in mouse and elephant metabolomes to determine whether fly-defined temporal pathway signatures showed evidence of mammalian age association (Fig. 5D). Several trajectory-linked pathways displayed age-related effects in one or both mammalian datasets, although the strength, direction, and completeness of these associations varied by species and pathway. Because age sampling strategies varied between species, concordance was evaluated by using the product of the overall direction of *Drosophila*-anchored metabolite abundances between the 5 day and 30 day timepoints and the mean species age effect coefficient from each species’ age-effect models (see Methods). Supporting analyses further summarized metabolite coverage and concordance within each *Drosophila*-defined trajectory cluster, clarifying where interpretation was constrained by pathway representation in the mammalian datasets (Supplementary Fig. S13).

Concordance analysis showed partial mammalian alignment with Drosophila trajectory-associated pathways, with strongest recurrent structure in lipid-associated and nucleotide-related categories. Some pathways aligned in direction, whereas others showed opposite mammalian directionality or limited metabolite coverage, emphasizing that pathway-level correspondence did not imply identical temporal behavior across species. Collectively, these analyses show that adult *Drosophila* aging is associated with extensive metabolomic remodeling that can be organized into distinct temporal trajectory classes. Although exact metabolite-level overlap across *Drosophila*, mouse, and elephant was limited, several *Drosophila*-defined age-associated pathways showed corresponding age-related structure in mammalian datasets. Carnitine/fatty acid metabolism, fatty acid classes, and nucleotide-related pathways emerged repeatedly across metabolite-level, pathway-level, score-based, and trajectory-concordance analyses. These results indicate that cross-species correspondence is more apparent at the level of metabolic pathways than individual metabolite identities, while the descriptive and exploratory nature of several pathway-overlap and score-based analyses warrants cautious interpretation. Supporting analyses further showed that these findings were robust to sex-specific evaluation, ranked-threshold sensitivity, pathway-overlap quantification, pathway-stratified scoring, directionality analysis, and concordance coverage assessment (Supplementary Figs. S1–S13).

## DISCUSSION

### Overview of cross-species age-associated metabolic remodeling

This study used a *Drosophila*-anchored comparative metabolomics framework to evaluate whether age-associated metabolic remodeling is shared across evolutionarily divergent species. The major finding is that age-associated metabolomic change was strongest and most temporally resolved in *Drosophila*, but several *Drosophila*-defined pathway signatures showed corresponding age-related structure in mouse and elephant. Exact metabolite-level overlap was limited, as expected given differences in organismal physiology, sample type, age range, and metabolite coverage. However, pathway-level overlap was broader, with recurrent signals in carnitine and fatty acid metabolism, monounsaturated and other fatty acid classes, nucleotide metabolism, indole/tryptophan metabolism, and central carbon metabolism. These results support a model in which aging is accompanied by partial pathway-level metabolic correspondence rather than strict overlap of individual metabolite changes.

### Drosophila as a temporal anchor for discovery of age-associated metabolism

Prior *Drosophila* metabolomics studies showed that the age-associated metabolome changes dynamically across adult life and can serve as a quantitative biomarker of age-related physiological state (Wang et al., 2022; Zhao et al., 2022). Our *Drosophila* dataset uniquely establishes temporal anchored support for broader age-associated metabolomic dynamics across species. Principal component analysis showed that adult age was the dominant axis of variation, and metabolite-level modeling identified widespread age-associated changes across the measured metabolome. These changes were not uniformly directional. Instead, age-associated metabolites resolved into distinct temporal classes, including progressive increases, progressive decreases, and transient midlife-associated patterns. This trajectory structure suggests that age-associated metabolism is not a simple monotonic deterioration of all pathways, but a dynamic process in which distinct metabolic programs remodel at different phases of adult aging.

### Fatty acid and acylcarnitine remodeling as a recurrent aging signal

Lipid-associated pathways were among the most biologically interpretable recurrent signals in this study. Rather than implying that identical lipid metabolites changed uniformly across species, these findings point to age-associated remodeling of lipid handling, fatty acid trafficking, and mitochondrial substrate use. Long-chain fatty acids require carnitine-dependent transport into mitochondria for β-oxidation, and acylcarnitines represent intermediates of fatty acid trafficking and mitochondrial substrate handling (Longo et al., 2016). In this context, age-associated changes in fatty acids and acylcarnitines may reflect an altered balance among fatty acid availability, mitochondrial oxidative capacity, and tissue energy demand.

This interpretation is consistent with human and experimental studies linking plasma acylcarnitine profiles to aging, incomplete fatty acid oxidation, insulin resistance, and altered metabolic flexibility (Adams et al., 2009; Jarrell et al., 2020; Koves et al., 2008). Aging and overnutrition can also compromise carnitine availability, mitochondrial performance, and metabolic control (Noland et al., 2009). Therefore, the lipid and acylcarnitine signal observed here is best interpreted as a pathway-level marker of altered mitochondrial lipid handling, rather than as evidence that the same lipid metabolites change identically across species. The present study does not establish that the observed mammalian lipid patterns are pathogenic, and the elephant lipid results should be considered exploratory given the cross-sectional and naturally heterogeneous sampling design.

Monounsaturated fatty acids may provide an additional window into age-associated metabolic state. Circulating monounsaturated fatty acids can reflect diet, adipose tissue composition, de novo lipogenesis, membrane remodeling, lipid mobilization, or altered substrate use. Prior metabolomics and Mendelian randomization analyses suggest that insulin resistance can alter circulating oleic and palmitoleic acid levels, linking these metabolites to metabolic disease pathways relevant to type 2 diabetes and cardiovascular disease (Mozaffarian et al., 2010; Nowak et al., 2016). Thus, age-associated monounsaturated fatty acid patterns in the mammalian datasets may reflect altered lipid partitioning, reduced metabolic flexibility, or compensatory lipid remodeling with age.

### Mitochondrial dysfunction and altered energy use

Age-associated lipid and acylcarnitine changes may reflect a mismatch between fatty acid availability, mitochondrial oxidative capacity, and tissue energy demand. Mitochondrial dysfunction is a core hallmark of aging and is closely connected to deregulated nutrient sensing, oxidative stress, chronic inflammation, and loss of proteostasis (López-Otín et al., 2023). Mitochondria also coordinate substrate oxidation, redox balance, and energetic adaptation across tissues, linking mitochondrial function to age-related metabolic disease (Amorim et al., 2022; Tan et al., 2025). In this context, the fatty acid and acylcarnitine signals observed here may represent systemic signatures of altered mitochondrial substrate handling rather than simple changes in lipid abundance.

### Nucleotide metabolism and purine turnover

The nucleotide-related signal suggests that age-associated metabolic remodeling also involves purine turnover, urate metabolism, and redox balance. This interpretation is supported by the appearance of purine-related metabolites such as xanthine and hypoxanthine among mammalian age-associated features. These metabolites are products of purine degradation and sit upstream of uric acid production. Hypoxanthine is oxidized to xanthine and then uric acid through xanthine oxidoreductase activity, a process that can generate reactive oxygen species depending on enzyme form and redox context (Battelli et al., 2016; Nelson & Voruganti, 2026). In humans, dysregulated purine metabolism is reported to be directly relevant to gout and hyperuricemia-associated conditions, including renal disease, metabolic syndrome, hypertension, diabetes, and cardiovascular disease (Du et al., 2024; Wang et al., 2019).

### Insights from long-lived mammalian species

The elephant dataset extends this framework beyond short-lived experimental models and provides an opportunity to ask whether age-associated metabolic themes are also detectable in long-lived mammalian physiology. Comparative metabolomics has shown that mammalian metabolomes are shaped by organ function, lineage, and longevity, suggesting that long-lived species may occupy metabolic states not fully represented by conventional laboratory models (Ma et al., 2015). The lipid- and nucleotide-related structure observed in elephants may reflect broad remodeling of mitochondrial fuel use, membrane and lipid handling, purine turnover, and redox-associated metabolism. Because the elephant dataset is cross-sectional and naturally heterogeneous, these patterns should be interpreted as supportive exploratory evidence that related metabolic systems vary with age in a long-lived mammalian context. Thus, the main contribution of the elephant dataset is to broaden the comparative scope of the study while reinforcing the distinction between pathway-level biological correspondence and strict conservation of metabolite identity, directionality, or timing.

### Limitations and future directions

Several limitations should be considered. First, the *Drosophila* dataset is the only dense adult time course; mouse is a young-versus-old contrast and elephant is cross-sectional. Therefore, mammalian data cannot validate the exact temporal shape of *Drosophila* trajectories. Instead, the mammalian analyses test whether *Drosophila*-defined pathways show age-associated directionality or pathway-level structure. Second, sample types differ across species. Whole-fly profiles integrate multiple tissues, whereas mouse and elephant plasma reflect circulating metabolites influenced by tissue release, uptake, diet, renal clearance, and systemic physiology. Third, pathway annotations differ in depth and specificity, and broad pathways such as fatty acid metabolism may include metabolites with distinct biochemical roles. Finally, the mammalian datasets demonstrate less statistical power than the *Drosophila* time course, so mammalian pathway-level findings should be interpreted as evidence of partial correspondence.

Future work should test whether the lipid and nucleotide pathways identified here are mechanistically linked to aging phenotypes. In *Drosophila*, targeted manipulation of fatty acid oxidation, carnitine transport, mitochondrial function, or nucleotide metabolism could determine whether these pathways modify lifespan, stress resistance, locomotor decline, or other age-associated traits. In mammals, larger longitudinal studies could determine whether acylcarnitine and monounsaturated fatty acid changes predict biological aging, frailty, insulin resistance, cardiovascular risk, or renal decline. Additional long-lived mammalian datasets would help determine whether the age-associated patterns in elephants represent general features of long-lived species or species-specific physiology.

### Conclusions

This study identifies recurrent age-associated remodeling of lipid, acylcarnitine, fatty acid, and nucleotide-related pathways across *Drosophila*, mouse, and elephant. The strongest cross-species signal is not strict redundance of individual metabolites, but partial pathway-level correspondence involving metabolic systems implicated in mitochondrial fuel use, lipid handling, purine turnover, and human age-related disease. These findings support a *Drosophila*-anchored comparative framework in which dense temporal metabolomics from a short-lived model organism can define age-associated pathway trajectories, while mammalian datasets test whether those pathways show broader relevance across lifespan scales and physiological contexts.

## Acknowledgements

We’d like to thank Game Rangers International, including Senior Research Officer Kasi Kalande and Research Assistant Lanos Chisaka, as well as African Parks and Zambia’s Department of National Parks and Wildlife for support of the study and assistance with elephant sample collection.

## Author Contributions

Computational analyses and manuscript writing completed by G.A.M. Elephant and mouse data organization provided by E.E.M. Fly data organization provided by S.R.S. Elephant plasma samples collected by I.O. and D.E.C. Targeted metabolomics was provided by T.N., A.D., and D.S. Mouse housing and plasma sampling completed by L.M.H. *Drosophila* samples and time course design provided by J.M.T. and T.C.K. Metabolomics coordination provided by J.M.T. Study conceptualization and computational guidance provided by H.T. Research group collaborative organization provided by D.E.C.

## REFERENCES

Adams, S. H., Hoppel, C. L., Lok, K. H., Zhao, L., Wong, S. W., Minkler, P. E., Garvey, W. T. (2009). Plasma acylcarnitine profiles suggest incomplete long-chain fatty acid β-oxidation and altered tricarboxylic acid cycle activity in type 2 diabetic African-American women. The Journal of Nutrition, 139, 1073–1081. 10.3945/jn.108.103754

Amorim, J. A., Coppotelli, G., Rolo, A. P., Palmeira, C. M., Ross, J. M., & Sinclair, D. A. (2022). Mitochondrial and metabolic dysfunction in ageing and age-related diseases. Nature Reviews Endocrinology, 18, 243–258. 10.1038/s41574-021-00626-7

Avanesov, A. S., Ma, S., Pierce, K. A., Yim, S. H., Lee, B. C., Clish, C. B., & Gladyshev, V. N. (2014). Age- and diet-associated metabolome remodeling characterizes the aging process driven by damage accumulation. eLife, 3, e02077. 10.7554/eLife.02077

Battelli, M. G., Polito, L., Bortolotti, M., & Bolognesi, A. (2016). Xanthine oxidoreductase in drug metabolism: Beyond a role as a detoxifying enzyme. Current Medicinal Chemistry, 23, 4027–4036. 10.2174/0929867323666160725091915

Bucaciuc Mracica, T., Anghel, A., Ion, C. F., Moraru, C. V., Tacutu, R., & Lazar, G. A. (2020). MetaboAge DB: A repository of known ageing-related changes in the human metabolome. Biogerontology, 21, 763–771. 10.1007/s10522-020-09892-w

Chusyd, D. E., Ackermans, N. L., Austad, S. N., Hof, P. R., Mielke, M. M., Sherwood, C. C., & Allison, D. B. (2021). Aging: What we can learn from elephants. Frontiers in Aging, 2, 726714. 10.3389/fragi.2021.726714

Di Francesco, A., Deighan, A. G., Litichevskiy, L., Chen, Z., Luciano, A., Robinson, L., Garland, G., Donato, H., Vincent, M., Schott, W., Wright, K. M., Raj, A., Prateek, G. V., Mullis, M., Hill, W. G., Zeidel, M. L., Peters, L. L., Harding, F., Botstein, D., … Churchill, G. A. (2024). Dietary restriction impacts health and lifespan of genetically diverse mice. Nature, 634(8034), 684–692. 10.1038/s41586-024-08026-3

Du, L., Zong, Y., Li, H., Wang, Q., Xie, L., Yang, B., Gao, J. (2024). Hyperuricemia and its related diseases: Mechanisms and advances in therapy. Signal Transduction and Targeted Therapy, 9, 212. 10.1038/s41392-024-01916-y

Dzieciatkowska, M., Issaian, A. V., Keele, G. R., Saviola, A., Stephenson, D., Bevers, S., Reisz, J. A., Haiman, Z. B., Nemkov, T., Fang, F., Moore, A. L., Deng, X., Stone, M., Kleinman, S., Norris, P. J., Wang, X., Thein, S.-L., Hod, E. A., Busch, M. P., D’Alessandro, A. (2026). A population-scale Red Blood Cell proteome atlas of 13,000 donors uncovers genetically encoded aging clocks predicting hemolysis, transfusion efficacy, and donor activity a decade later (p. 2026.03.07.710284). bioRxiv. 10.64898/2026.03.07.710284

Evangelakou, Z., Manola, M., Gumeni, S., & Trougakos, I. P. (2019). Nutrigenomics as a tool to study the impact of diet on aging and age-related diseases: The Drosophila approach. Genes & Nutrition, 14, 12. 10.1186/s12263-019-0638-6

Hanks, J. (1972). Growth of the African elephant (Loxodonta africana). East African Wildlife Journal, 10(4), 251–272. 10.1111/j.1365-2028.1972.tb00870.x

Hoffman, J. M., Soltow, Q. A., Li, S., Sidik, A., Jones, D. P., & Promislow, D. E. L. (2014). Effects of age, sex, and genotype on high-sensitivity metabolomic profiles in the fruit fly, Drosophila melanogaster. Aging Cell, 13(4), 596–604. 10.1111/acel.12215

Holsopple, J. M., Smoot, S. R., Popodi, E. M., Colbourne, J. K., Shaw, J. R., Oliver, B., Kaufman, T. C., & Tennessen, J. M. (2023). Assessment of chemical toxicity in adult Drosophila melanogaster. Journal of Visualized Experiments, 193, e65029. 10.3791/65029

Holtze, S., Gorshkova, E., Braude, S., Cellerino, A., Dammann, P., Hildebrandt, T. B., Sahm, A. (2021). Alternative animal models of aging research. Frontiers in Molecular Biosciences, 8, 660959. 10.3389/fmolb.2021.660959

Huang, H., Chen, Y., Xu, W., Cao, L., Qian, K., Bischof, E., Kennedy, B. K., & Pu, J. (2025). Decoding aging clocks: New insights from metabolomics. Cell Metabolism, 37, 34–58. 10.1016/j.cmet.2024.11.007

Ibáñez de Opakua, A., Conde, R., de Diego, A., Bizkarguenaga, M., Embade, N., Lu, S. C., Mato, J. M., & Millet, O. (2025). Metabolomic-based aging clocks. npj Metabolic Health and Disease, 3, 35. 10.1038/s44324-025-00078-x

Jankowski, C. S. R., Samarah, L. Z., MacArthur, M. R., Mitchell, S. J., Weilandt, D. R., Hunter, C. J., … Rabinowitz, J. D. (2025). Aged mice exhibit widespread metabolic changes but preserved major fluxes. Cell Metabolism, 37, 2280–2294.e4. 10.1016/j.cmet.2025.09.009

Jarrell, Z. R., Smith, M. R., Hu, X., Orr, M., Liu, K. H., Quyyumi, A. A., Jones, D. P., & Go, Y.-M. (2020). Plasma acylcarnitine levels increase with healthy aging. Aging, 12, 13555–13570. 10.18632/aging.103462

Johnson, L. C., Parker, K., Aguirre, B. F., Nemkov, T. G., D’Alessandro, A., Johnson, S. A., Seals, D. R., & Martens, C. R. (2019). The plasma metabolome as a predictor of biological aging in humans. GeroScience, 41(6), 895–906. 10.1007/s11357-019-00123-w

Koves, T. R., Ussher, J. R., Noland, R. C., Slentz, D., Mosedale, M., Ilkayeva, O., Muoio, D. M. (2008). Mitochondrial overload and incomplete fatty acid oxidation contribute to skeletal muscle insulin resistance. Cell Metabolism, 7, 45–56. 10.1016/j.cmet.2007.10.013

Kuiper, L. M., Polinder-Bos, H. A., Bizzarri, D., Vojinovic, D., Vallerga, C. L., Dollé, M. E. T., van Meurs, J. B. J. (2023). Epigenetic and metabolomic biomarkers for biological age: A comparative analysis of mortality and frailty risk. The Journals of Gerontology: Series A, 78, 1753–1762. 10.1093/gerona/glad137

Longo, N., Frigeni, M., & Pasquali, M. (2016). Carnitine transport and fatty acid oxidation. Biochimica et Biophysica Acta, 1863, 2422–2435. 10.1016/j.bbamcr.2016.01.023

López-Otín, C., Blasco, M. A., Partridge, L., Serrano, M., & Kroemer, G. (2023). Hallmarks of aging: An expanding universe. Cell, 186, 243–278. 10.1016/j.cell.2022.11.001

Ma, S., & Gladyshev, V. N. (2017). Molecular signatures of longevity: Insights from cross-species comparative studies. Seminars in Cell & Developmental Biology, 70, 190–203. 10.1016/j.semcdb.2017.08.007

Ma, S., Yim, S. H., Lee, S. G., Kim, E. B., Lee, S. R., Chang, K. T., Buffenstein, R., Lewis, K. N., Park, T. J., Miller, R. A., Clish, C. B., & Gladyshev, V. N. (2015). Organization of the mammalian metabolome according to organ function, lineage specialization, and longevity. Cell Metabolism, 22, 332–343. 10.1016/j.cmet.2015.07.005

Menni, C., Kastenmüller, G., Petersen, A. K., Bell, J. T., Psatha, M., Tsai, P.-C., Valdes, A. M. (2013). Metabolomic markers reveal novel pathways of ageing and early development in human populations. International Journal of Epidemiology, 42, 1111–1119. 10.1093/ije/dyt094

Moon, S. J., Hu, Y., Dzieciatkowska, M., Kim, A.-R., Asara, J. M., D’Alessandro, A., & Perrimon, N. (2026). Modeling tissue-specific Drosophila metabolism identifies high sugar diet-induced metabolic dysregulation in muscle at reaction and pathway levels. Nature Communications, 17, 1692. 10.1038/s41467-026-68395-3

Mozaffarian, D., Cao, H., King, I. B., Lemaitre, R. N., Song, X., Siscovick, D. S., & Hotamisligil, G. S. (2010). Circulating palmitoleic acid and risk of metabolic abnormalities and new-onset diabetes. The American Journal of Clinical Nutrition, 92, 1350–1358. 10.3945/ajcn.110.003970

Mutz, J., Iniesta, R., & Lewis, C. M. (2024). Metabolomic age (MileAge) predicts health and life span: A comparison of multiple machine learning algorithms. Science Advances, 10, eadp3743. 10.1126/sciadv.adp3743

Nelson, K. L., & Voruganti, V. S. (2026). Implication of xanthine oxidoreductase in oxidative stress-related chronic diseases. Frontiers in Endocrinology, 16, 1662037. 10.3389/fendo.2025.1662037

Nemkov, T., Reisz, J. A., Gehrke, S., Hansen, K. C., & D’Alessandro, A. (2019). High-throughput metabolomics: Isocratic and gradient mass spectrometry-based methods. In A. D’Alessandro (Ed.), High-throughput metabolomics (Methods in Molecular Biology, Vol. 1978, pp. 13–26). Humana. 10.1007/978-1-4939-9236-2_2

Noland, R. C., Koves, T. R., Seiler, S. E., Lum, H., Lust, R. M., Ilkayeva, O., Stevens, R. D., Hegardt, F. G., & Muoio, D. M. (2009). Carnitine insufficiency caused by aging and overnutrition compromises mitochondrial performance and metabolic control. Journal of Biological Chemistry, 284, 22840–22852. 10.1074/jbc.M109.032888

Nowak, C., Salihovic, S., Ganna, A., Brandmaier, S., Tukiainen, T., Broeckling, C. D., Fall, T. (2016). Effect of insulin resistance on monounsaturated fatty acid levels: A multi-cohort non-targeted metabolomics and Mendelian randomization study. PLOS Genetics, 12, e1006379. 10.1371/journal.pgen.1006379

Oh, H. S.-H., Rutledge, J., Nachun, D., Pálovics, R., Abiose, O., Moran-Losada, P., Wyss-Coray, T. (2023). Organ aging signatures in the plasma proteome track health and disease. Nature, 624, 164–172. 10.1038/s41586-023-06802-1

Fleck SA, Biswas P, DeWitt ED, Knuteson RL, Eisman RC, Nemkov T, D’Alessandro A, Tennessen JM, Rideout E, Weaver LN. Auxin exposure disrupts feeding behavior and fatty acid metabolism in adult Drosophila. Elife. 2024 Jan 19;12:RP91953. doi: 10.7554/eLife.91953. PMID: 38240746; PMCID: PMC10945601. https://doi.org/10.7554/eLife.91953.3

Öztürk-Çolak, A., Marygold, S. J., Antonazzo, G., Attrill, H., Goutte-Gattat, D., Jenkins, V. K., FlyBase Consortium. (2024). FlyBase: Updates to the Drosophila genes and genomes database. Genetics, 227, iyad211. 10.1093/genetics/iyad211

Piper, M. D. W., & Partridge, L. (2018). Drosophila as a model for ageing. Biochimica Et Biophysica Acta. Molecular Basis of Disease, 1864(9 Pt A), 2707–2717. 10.1016/j.bbadis.2017.09.016

Reisz, J. A., Earley, E. J., Nemkov, T., Key, A., Stephenson, D., Keele, G. R., Dzieciatkowska, M., Spitalnik, S. L., Hod, E. A., Kleinman, S., Roubinian, N. H., Gladwin, M. T., Hansen, K. C., Norris, P. J., Busch, M. P., Zimring, J. C., Churchill, G. A., Page, G. P., & D’Alessandro, A. (2025). Arginine metabolism is a biomarker of red blood cell and human aging. Aging Cell, 24(2), e14388. 10.1111/acel.14388

Sebastiani, P., Monti, S., Lustgarten, M., Song, Z., Ellis, D., Tian, Q., Long Life Family Study Research Group. (2024). Metabolite signatures of chronological age, aging, survival, and longevity. Cell Reports, 43, 114913. 10.1016/j.celrep.2024.114913

Tan, S., Carbone, M. A., Zhou, S., Morozova, T., Arya, G., Zhang, F., Anholt, R. R. H., Huang, W., & Mackay, T. F. C. (2025). Effects of aging on gene expression networks in the Drosophila genetic reference panel. Cell Reports, 44(7), 115999. 10.1016/j.celrep.2025.115999

van Dam, E., van Leeuwen, L. A. G., dos Santos, E., James, J., Best, L., Lennicke, C., Cochemé, H. M. (2020). Sugar-induced obesity and insulin resistance are uncoupled from shortened survival in Drosophila. Cell Metabolism, 31, 710–725.e7. 10.1016/j.cmet.2020.02.016

Vecchié, D., Anholt, R. R. H., Mackay, T. F. C., & De Luca, M. (2026). Selection for Postponed Senescence in Drosophila melanogaster Reveals Distinct Metabolic Aging Trajectories Modifiable by the Angiotensin-Converting Enzyme Inhibitor Lisinopril. Aging Cell, 25(2), e70375. 10.1111/acel.70375

Wang, R., Yin, Y., Li, J., Wang, H., Lv, W., Gao, Y., Zhu, Z.-J. (2022). Global stable-isotope tracing metabolomics reveals system-wide metabolic alterations in aging Drosophila. Nature Communications, 13, 3518. 10.1038/s41467-022-31268-6

Wang, Y., Deng, M., Deng, B., Ye, L., Fei, X., & Huang, Z. (2019). Study on the diagnosis of gout with xanthine and hypoxanthine. Journal of Clinical Laboratory Analysis, 33, e22868. 10.1002/jcla.22868

Zhao, X., Golic, F. T., Harrison, B. R., Manoj, M., Hoffman, E. V., Simon, N., Promislow, D. E. L. (2022). The metabolome as a biomarker of aging in Drosophila melanogaster. Aging Cell, 21, e13548. 10.1111/acel.13548

