## Supporting Information for "Comparative metabolomics identifies recurrent age-associated pathway remodeling across species"

The Supporting Information contains supplemental analyses that provide additional context for the primary findings presented in Figures 1–5. These analyses examine sex-specific metabolomic structure, species-specific age-associated metabolites, pathway annotation and overlap, ranked-threshold sensitivity, pathway-level aging scores, directionality of age effects, and cross-species concordance of *Drosophila*-defined trajectory classes. Collectively, the supplemental figures support the central conclusion that aging-associated metabolomic remodeling is more consistently observed at the pathway level than at the level of identical metabolite changes across species.

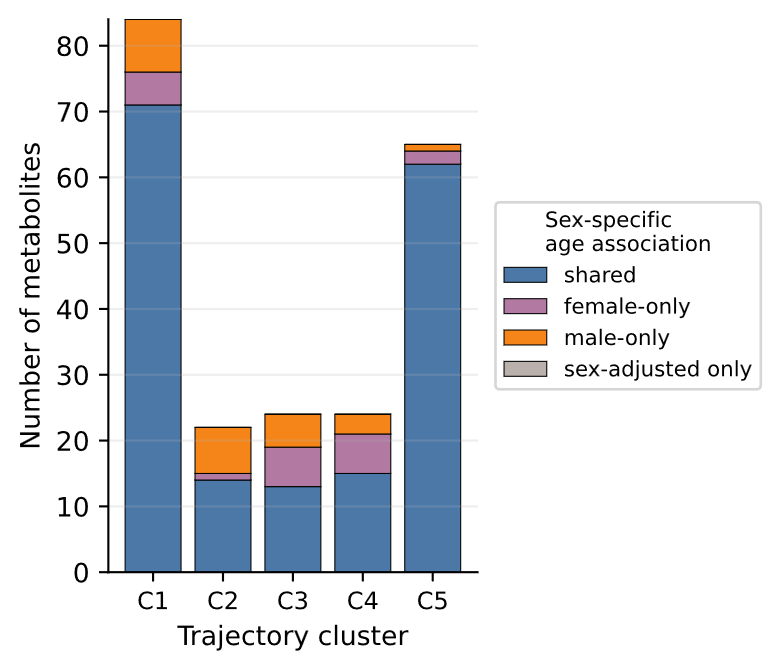

Supplementary Figure 1: Supplementary Figure 1. Sex-specific age-associated metabolite counts within Drosophila trajectory clusters

We deconstruct the Drosophila trajectory analysis shown in Figure 5 by summarizing how age-associated metabolites are distributed between females and males within each Drosophila temporal cluster. This analysis evaluates whether the trajectory clusters are dominated by one sex or whether both sexes contribute to the age-associated metabolite patterns. By stratifying metabolite counts by sex and cluster, the figure provides additional context for interpreting the temporal classes as aging-associated patterns rather than artifacts of sex imbalance. This supplemental analysis supports the main conclusion that Drosophila age-associated metabolites organize into distinct temporal trajectories, while also showing whether those trajectories contain sex-specific structure.

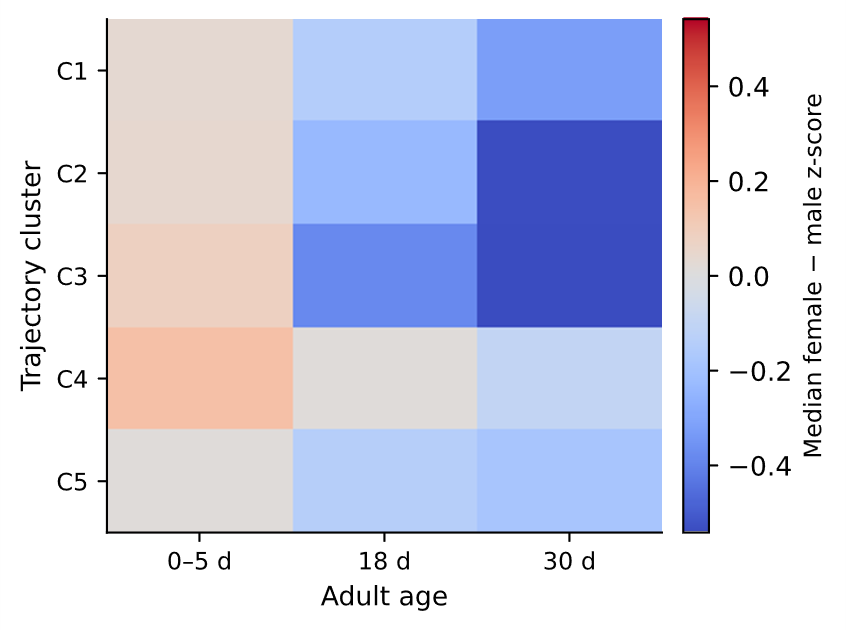

Supplementary Figure 2. Median female-minus-male z-score differences within age-associated metabolite clusters

To support the interpretation of sex-related structure within the Drosophila aging trajectories, we summarize metabolite abundances as the median female-minus-male difference in z-scored abundance within each trajectory cluster. This analysis shows whether females and males differ systematically in metabolite abundance within the same age-associated trajectory classes. The figure complements Figure 5 by separating temporal aging patterns from sex-associated abundance differences, helping to determine whether individual clusters reflect broadly shared age trajectories or contain consistent sex-biased metabolic features.

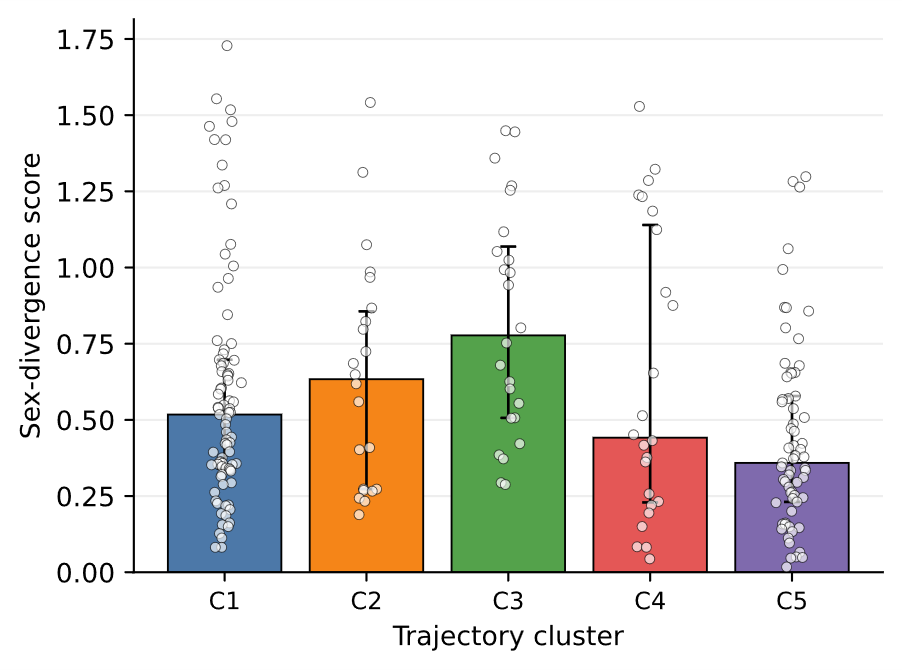

Supplementary Figure 3. Root mean-squared sex-divergence score within Drosophila temporal clusters

We provide a quantitative summary of sex divergence across Drosophila age-associated trajectory clusters. For each metabolite, female-minus-male mean z-score differences were calculated at each adult age, and the root mean square of these age-specific differences was used to summarize the magnitude of sex divergence. Cluster-level summaries then describe whether some temporal trajectory classes show stronger sex-dependent separation than others. This figure supports Figure 5 by showing that Drosophila aging trajectories can be evaluated not only by their temporal direction but also by the degree to which male and female metabolite profiles diverge within those trajectories.

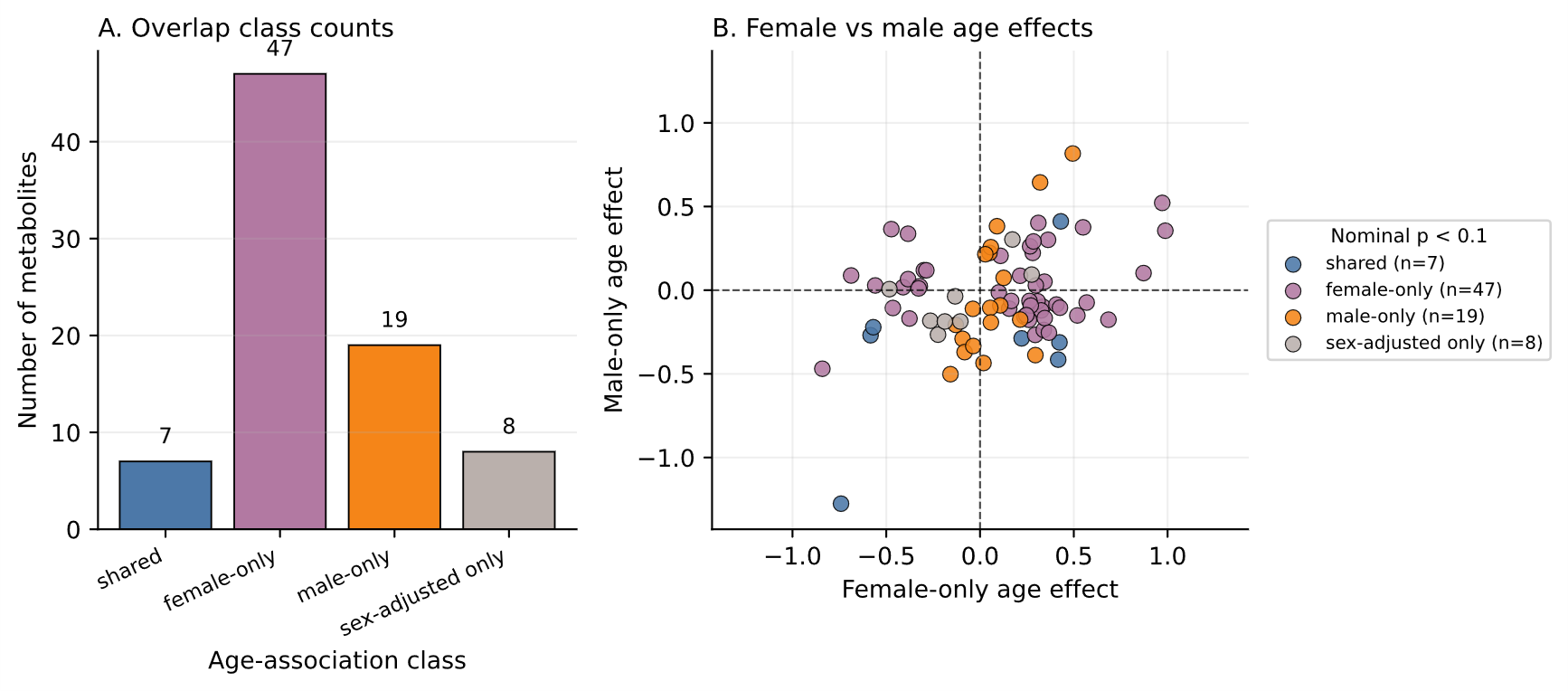

Supplementary Figure 4. Elephant sex-specific age-associated metabolite quantity and effect

The elephant age-effect analysis shown in Figure 3 is expanded by evaluating whether age-associated metabolite signals differ by sex. Because the elephant dataset is cross-sectional and includes both female and male individuals across a natural age gradient, sex-specific visualization provides important context for interpreting age effects. This supplemental figure helps distinguish metabolites that show broadly age-associated behavior from those whose apparent age association may be influenced by sex-specific abundance or effect patterns. It therefore supports cautious interpretation of elephant age-associated metabolites in the main text.

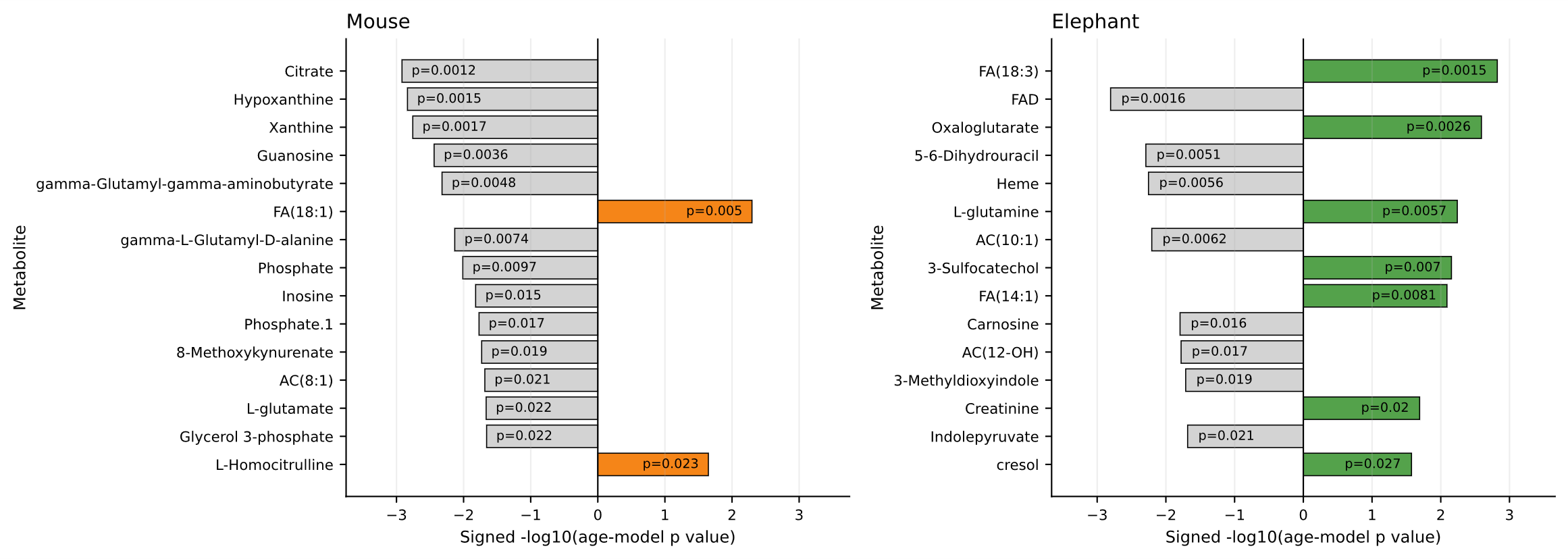

Supplementary Figure 5. Significant age-effect metabolites per species

Age-effect metabolites per species supports Figures 2 and 3 by summarizing the number and identity of metabolites associated with age in each species. Drosophila, mouse, and elephant differ in sample type, sampling density, age structure, and statistical power; therefore, this supplemental figure provides a direct comparison of metabolite-level age-effect output across datasets (p<0.1). The figure supports the main conclusion that Drosophila shows the strongest and most extensive age-associated metabolomic remodeling, whereas mouse and elephant show more limited metabolite-level signals but still retain pathway-level age-associated structure.

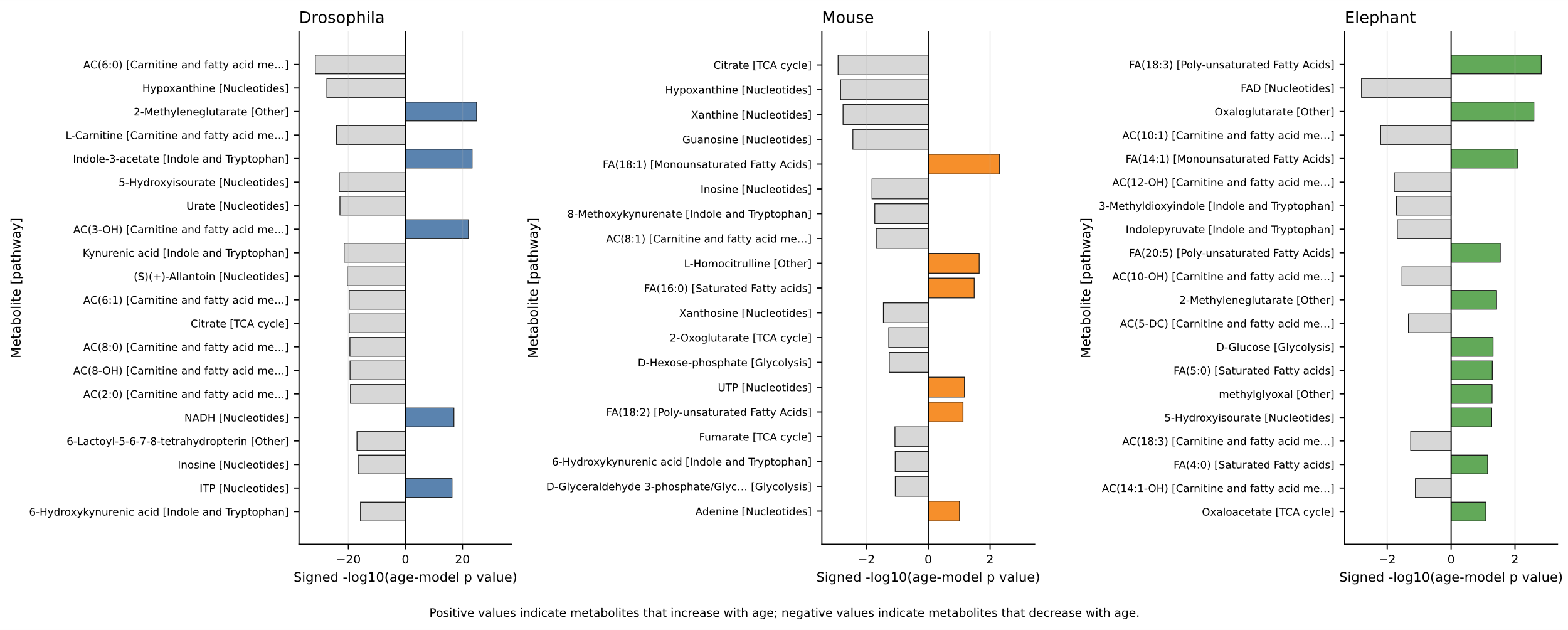

Supplementary Figure 6. Significant age-associated metabolites across species and mapped pathways

We support the pathway-overlap analyses shown in Figures 3 and 4 by connecting significant age-associated metabolites to their annotated pathways (p<0.1). This figure provides the metabolite-level basis for pathway-level interpretation, showing which metabolites contribute to each represented metabolic category across species. It helps clarify that pathway overlap does not necessarily imply identical metabolite changes across Drosophila, mouse, and elephant. Instead, it shows how different age-associated metabolites can map to related pathway categories, supporting the manuscript’s pathway-centered comparative framework.

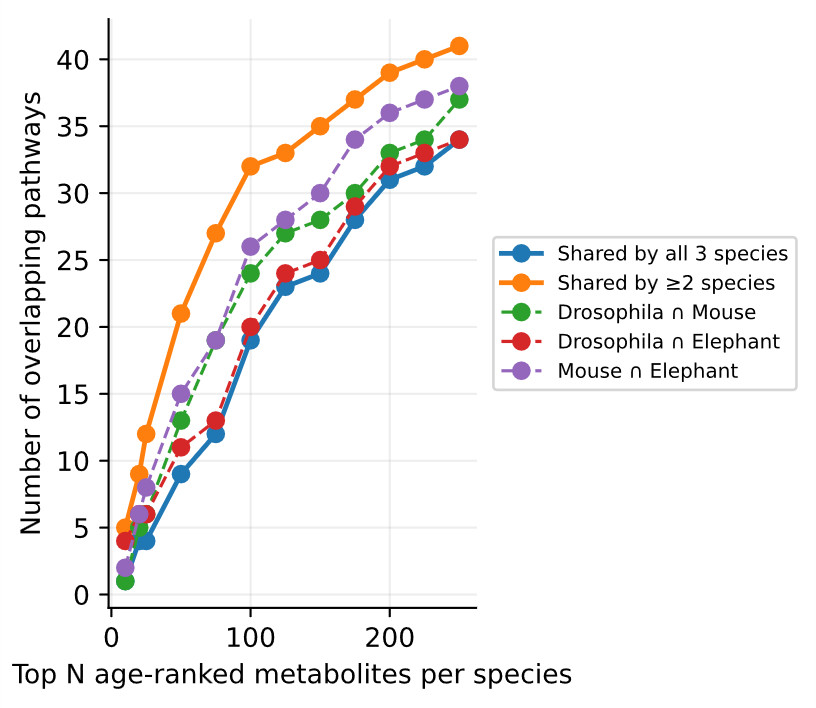

Supplementary Figure 7. Ranked age-associated metabolite pathway sensitivity analysis per species

To support the ranked pathway-overlap analysis used to compare species with different statistical power, we provide a sensitivity analysis using different topN age-ranked metabolites per species and their intersections. Because the mouse and elephant datasets contain fewer samples and less dense age sampling than the Drosophila time course, strict significance thresholds may underestimate shared pathway structure. This sensitivity analysis evaluates how pathway representation changes when age-associated metabolites are ranked by statistical evidence rather than only by hard significance cutoffs. The figure supports the main pathway-overlap conclusions by showing whether recurrent pathways remain evident across alternative ranked-feature thresholds. The null hypothesis was not formally evaluated for this descriptive ranked-overlap analysis; the analysis was intended to assess whether pathway representation was stable across alternative ranked feature counts.

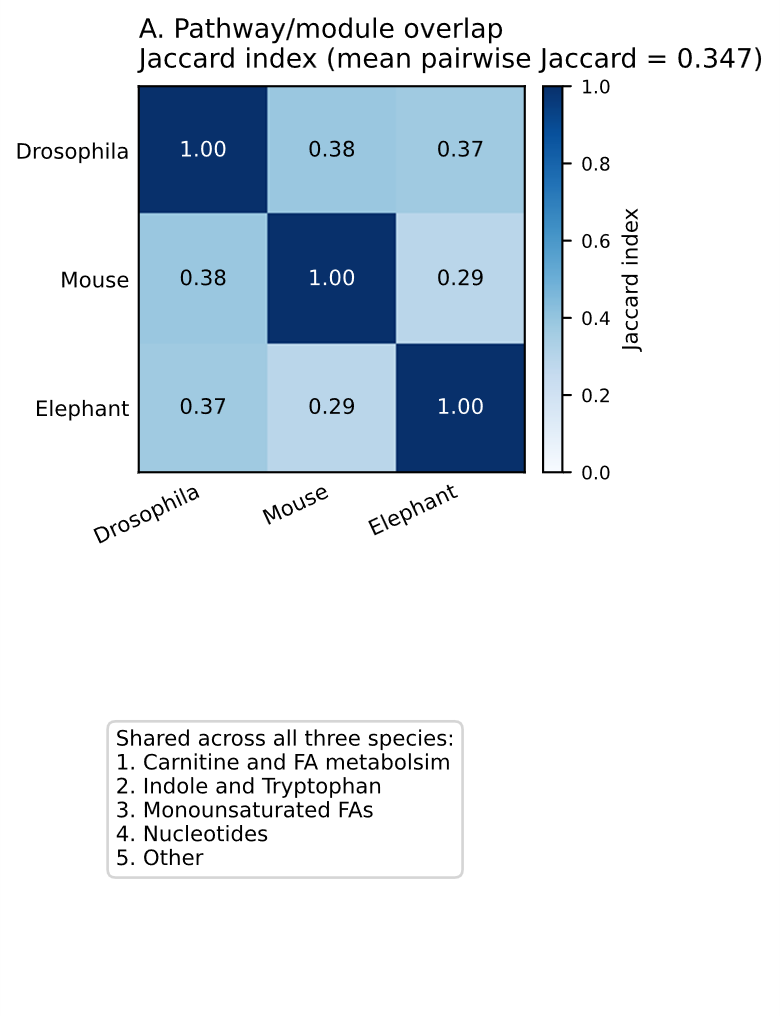

Supplementary Figure 8. Jaccard similarity of whole aging-associated pathway profiles between species

In this figure we quantify the similarity between species using Jaccard indices. Rather than focusing on individual metabolites, this analysis compares the overlap of aging-associated pathway profiles across Drosophila, mouse, and elephant. The figure provides a compact measure of pathway-level similarity and supports the conclusion that cross-species correspondence is stronger at the pathway level than at the individual-metabolite level. It also provides a quantitative basis for describing partial pathway-level correspondence.

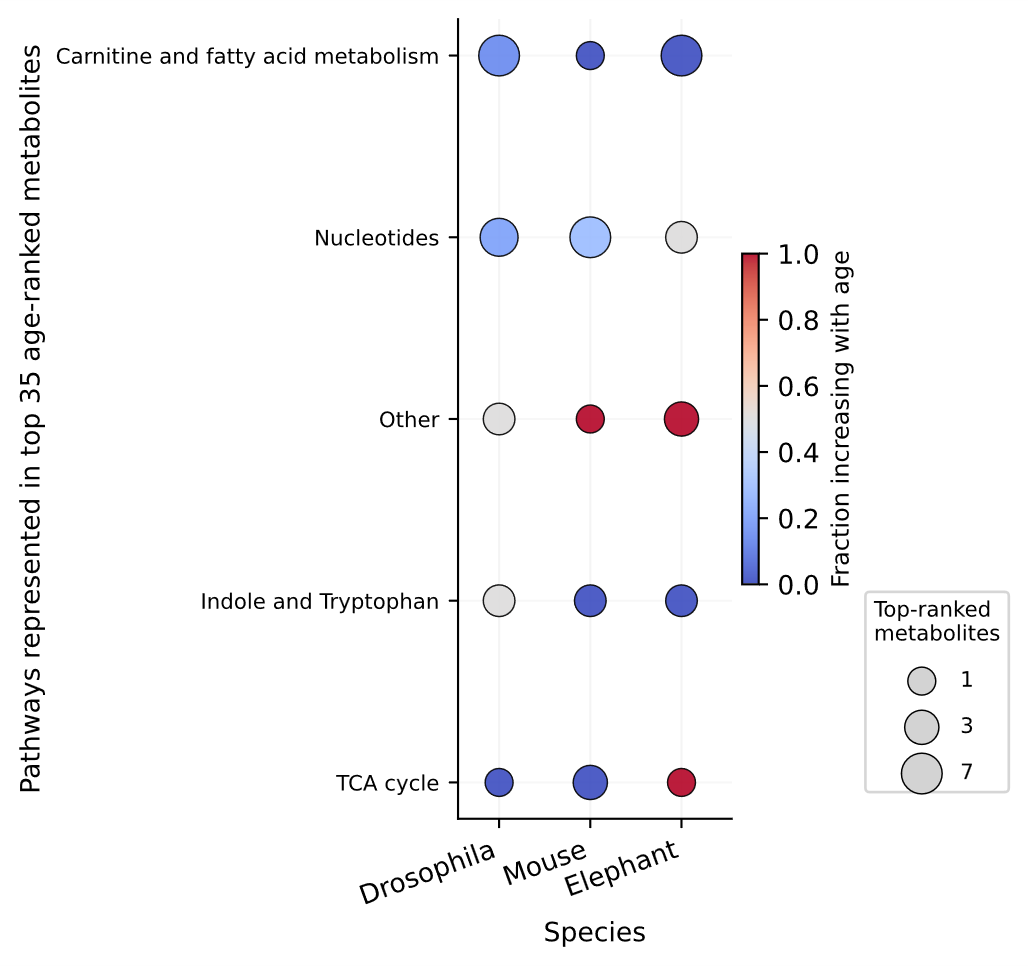

Supplementary Figure 9. Prevalence of accumulating age-associated metabolites per represented pathway

We elaborate the interpretation of pathway directionality by showing the proportion of age-associated metabolites within each pathway that increase with age. This analysis is important because pathways can contain metabolites with mixed age-effect directions. By summarizing the prevalence of accumulating metabolites, the figure helps distinguish pathways characterized primarily by age-associated accumulation from those showing mixed or decreasing metabolite behavior. This supports interpretation of lipid, fatty acid, nucleotide, and related metabolic pathways in the main Results and Discussion.

| **Pathway** | **Drosophila age-associated metabolites defining pathway** | **Drosophila metabolites used** | **Mouse metabolites used** | **Elephant metabolites used** |
| --- | --- | --- | --- | --- |
| Aminosugars | 6 | 6 | 5 | 4 |
| Arginine and proline metabolism | 4 | 4 | 10 | 8 |
| Carnitine and fatty acid metabolism | 28 | 30 | 36 | 32 |
| GSH homeostasis | 6 | 7 | 6 | 3 |
| Glycerophospholipid biosynthesis | 7 | 7 | 4 | 2 |
| Glycolysis | 2 | 4 | 6 | 4 |
| Indole and Tryptophan | 9 | 11 | 12 | 12 |
| Monounsaturated Fatty Acids | 3 | 4 | 4 | 4 |
| Nucleotides | 38 | 43 | 29 | 25 |
| Other | 8 | 10 | 17 | 16 |
| Panthothenate metabolism | 4 | 4 | 3 | 2 |
| Poly-unsaturated Fatty Acids | 4 | 4 | 6 | 6 |
| Proteolysis | 4 | 4 | 4 | 4 |
| Saturated Fatty acids | 7 | 9 | 9 | 9 |
| Serine biosynthesis and one-carbon metabolism | 7 | 7 | 4 | 4 |
| Sulfur metabolism | 3 | 3 | 3 | 3 |
| TCA cycle | 6 | 6 | 5 | 6 |
| Tyrosine Metabolism | 4 | 6 | 8 | 7 |
| Urea cycle | 2 | 2 | 2 | 2 |

Supplementary Table 1. Drosophila anchored age-associated metabolite pathways defining global aging score across species.

Pathways were defined from FDR-significant age-associated metabolites in *Drosophila*. For each retained pathway, pathway aging scores were calculated separately within each species using the metabolites detected and modeled in that species and assigned to the corresponding pathway. Pathways were retained for cross-species comparison when at least two metabolites were available for scoring in each species.

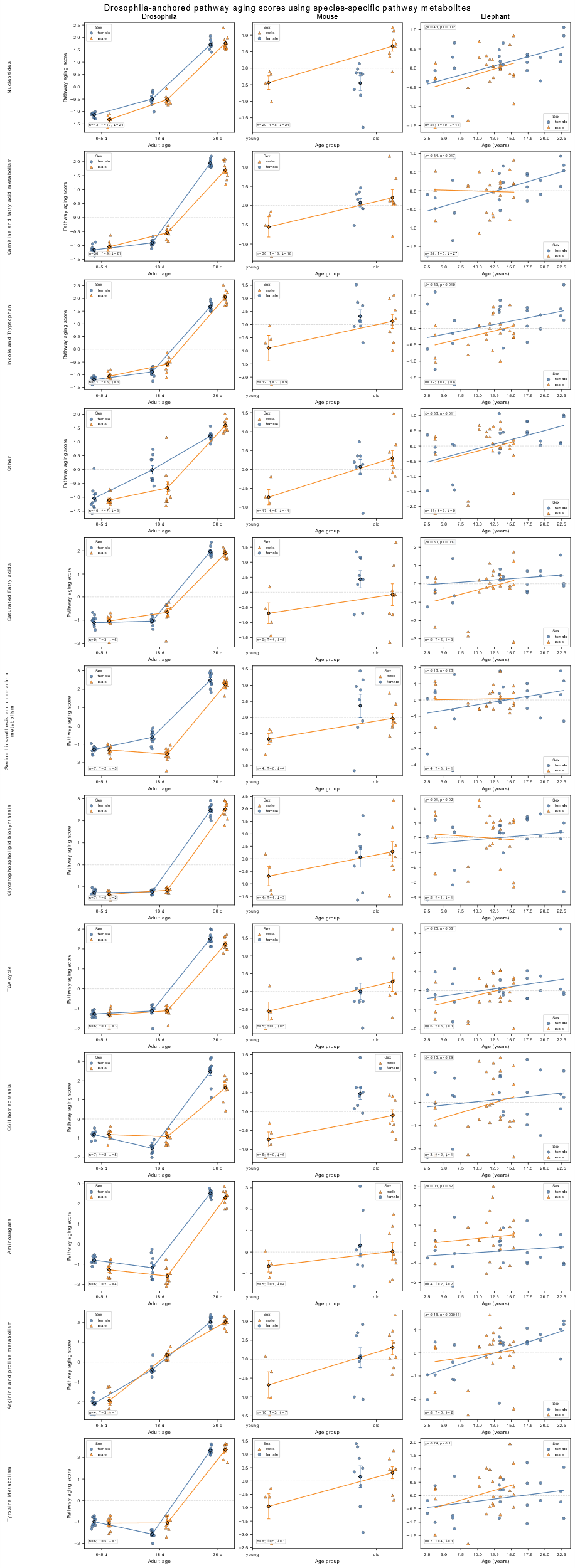

Supplementary Figure 10. Pathway-stratified metabolic aging scores for each species

We stratify the global pathway-aging score analysis shown in Figure 4 by pathway used to compute the score. Whereas the main figure summarizes pathway-anchored metabolic aging scores across species, this supplemental figure separates the score by individual pathway. This allows identification of which Drosophila-defined aging-associated pathways contribute most strongly to age-related score structure within Drosophila, mouse, and elephant. The figure supports the conclusion that the global aging score reflects coordinated pathway-level remodeling rather than a single dominant metabolite or pathway.

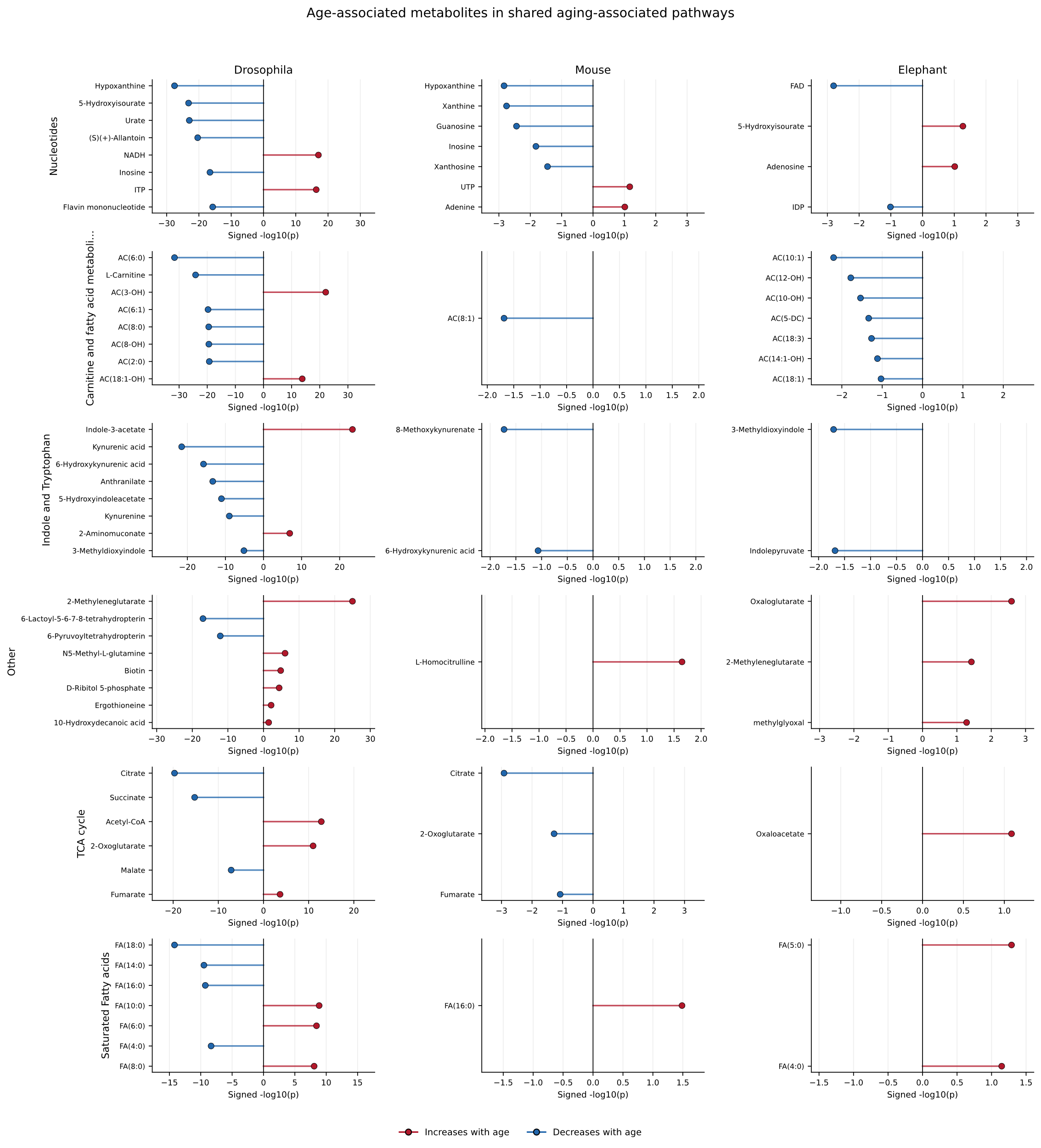

Supplementary Figure 11. Pathway-stratified age-associated metabolite directionality per species

To bolster the interpretation of species-specific directionality in the pathway-aging score and concordance analyses, we stratify directional analysis by age-associated pathway. For each species, age-associated metabolites within each pathway are summarized according to whether they increase or decrease with age. This figure helps clarify why the same pathway can appear in multiple species while showing different directionality or metabolite composition. It supports the manuscript’s central distinction between pathway-level correspondence and strict overlap of individual metabolite behavior.

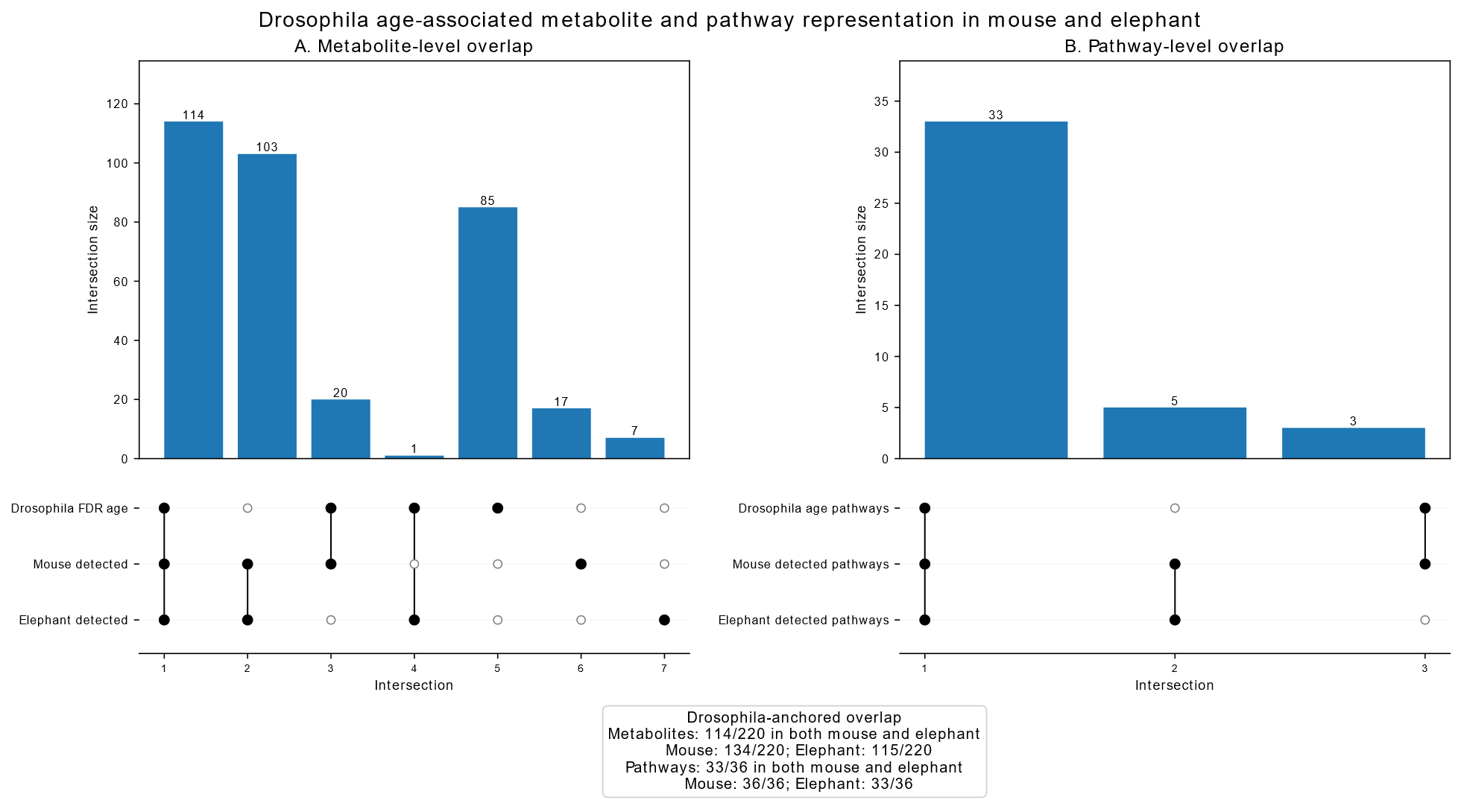

Supplementary Figure 12. UpSet plots comparing metabolite and pathway overlap structure across species

Supplementary Figure 12 supports the overlap analyses shown in Figures 3 and 4 by providing an alternative visualization of cross-species overlap. The UpSet plots summarize intersection patterns for metabolites and pathways across Drosophila, mouse, and elephant. This representation is useful because it shows both shared and species-specific sets without relying on Venn diagrams alone. The figure reinforces the main finding that exact metabolite-level overlap is limited, whereas pathway-level overlap is broader and more informative for cross-species comparison.

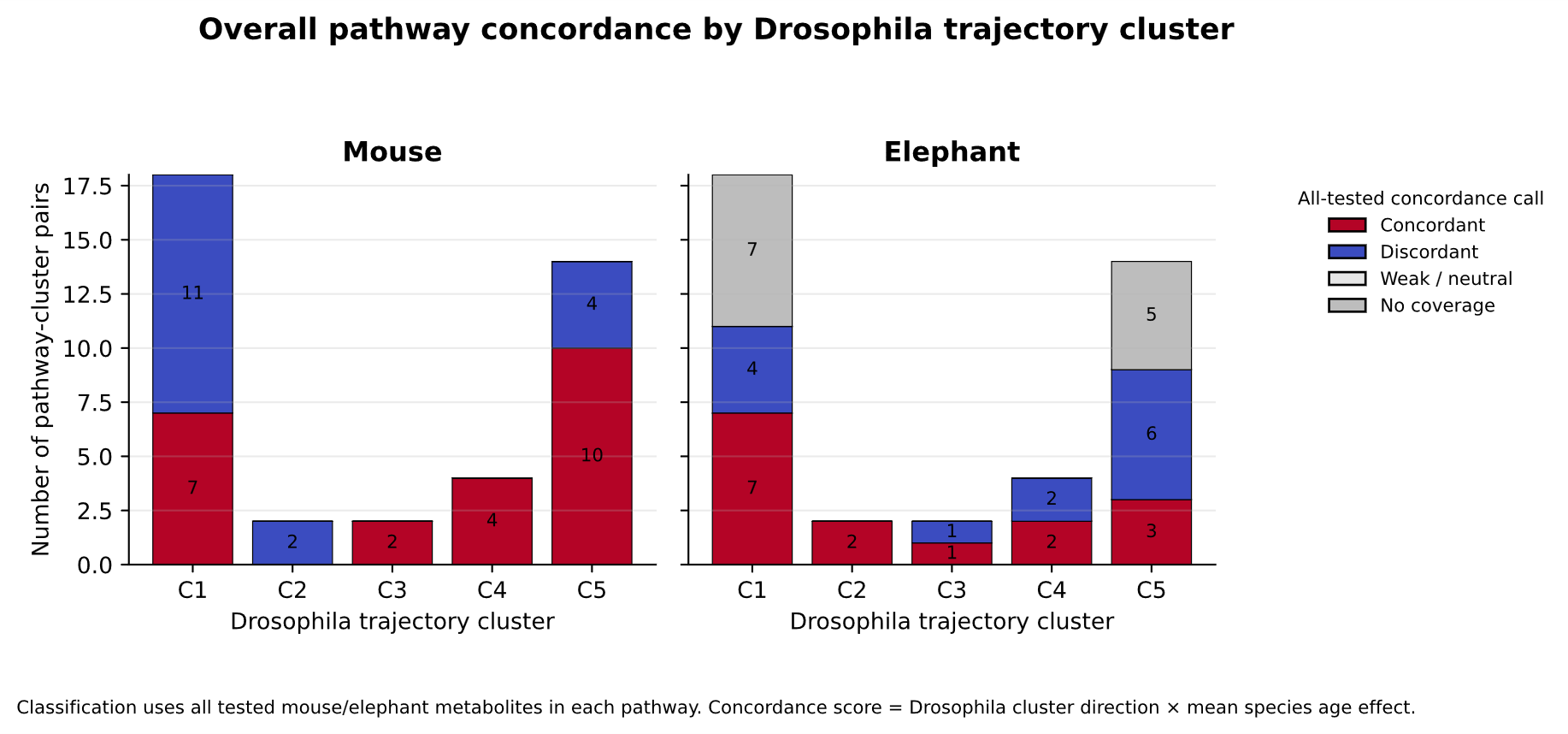

Supplementary Figure 13. Coverage and concordance of metabolites within each Drosophila-defined trajectory cluster

We simplify the trajectory-concordance analysis shown in Figure 5 by showing species-specific stacked bar plots of concordance magnitude by trajectory cluster. For each Drosophila trajectory cluster, this figure summarizes how well the corresponding pathway-associated metabolites are represented in mouse and elephant and whether mammalian age effects align with the Drosophila trajectory direction. This is important because concordance scores depend on both pathway coverage and directionality. The figure therefore provides supporting evidence for interpreting mammalian correspondence to Drosophila-defined temporal aging programs, while also identifying cases where limited metabolite coverage constrains interpretation.
